# Towards patient-specific biomechanical human brain models

**DOI:** 10.64898/2026.04.16.718870

**Authors:** Nicole Tueni, Benedikt Rauh, Jan Hinrichsen, Stefan Rampp, Arnd Doerfler, Silvia Budday

## Abstract

Reliable characterization of spatial variations in brain tissue stiffness is essential for predictive biomechanical modeling, yet most current methods rely on coarse regional parameter assignments based on invasive mechanical testing. In this study, we propose a new approach to obtain subject-specific mechanical properties at voxel resolution from *in vivo* diffusion tensor magnetic resonance imaging (DTI) based on a linear regression between the fractional anisotropy (FA) from DTI and experimentally measured stiffness values. To assess how such heterogeneity in mechanical properties influences simulated brain deformation, we construct a finite element model based on two material parameterizations of the same human brain: one employing nine anatomically defined regions, each with uniform material parameters, and another in which the shear modulus is assigned voxel-wise on the corresponding FA value. Applying this FA-stiffness mapping yields a smoothly varying mechanical property distribution that better captures local microstructural differences not represented by region-wise parameterizations. Both parameterizations are subjected to an identical atrophy-driven loading scenario. They exhibit comparable overall volume loss, but diverge in regional behavior. The voxel-resolved parameterization predicts more pronounced ventricular expansion and differs in the displacement and stretch distributions, indicating that variability in stiffness can alter local predicted responses even when global outcomes appear similar.

This work presents a pipeline for estimating individualized mechanical properties directly from imaging protocols that are routinely performed on patients, with important implications for brain biomechanics. While the approach depends on a simplified linear FA-stiffness relation and assumes isotropic constitutive behavior, it provides a framework for integrating imaging-based microstructure into subject-specific simulations. Future validation against *in vivo* or experimental deformation data is needed to determine the fidelity and clinical utility of FA-derived stiffness fields.

## 1. Introduction

Brain biomechanics has emerged as a crucial field of research with significant clinical relevance, particularly in neurosurgery, neurodegeneration, and brain injury assessment. By simulating patient-specific biomechanical responses, computational brain models support surgical planning, improve neuronavigation accuracy, and improve predictive tools for trauma and disease progression. Such models hold substantial promise for the advancement of diagnosis, personalized therapies, and improved clinical decision making in neurological care [1, 2, 3].

A major challenge for the successful application of computational brain models in clinical workflows is that such models must realistically represent the complexity of the brain, ideally in its *in vivo* state. A particularly significant advancement in this direction is the move away from averaged large-strain parameters obtained from *ex vivo* experiments toward patient-specific, locally varying parameter distributions that can be estimated from *in vivo* data, allowing the tissue behavior to be captured at the individual level.

Over the past decades, brain biomechanical modeling has undergone substantial development, both in terms of geometry representation and constitutive modeling. Early models relied on simplified geometries, such as fluid-filled two-dimensional spheres [4], before transitioning to three-dimensional models that progressively incorporated realistic anatomical structure [5]. Subsequent work assigned different mechanical properties to white and gray matter [6], and later included detailed cortical folding patterns with gyri and sulci [7]. Magnetic resonance imaging (MRI) made it possible to extract accurate patient-specific brain geometries and generate voxel-wise meshes with distinct anatomical regions [8, 9, 10, 11]. More recent efforts have moved toward comprehensive digital twins that integrate multimodal data, such as MRI and EEG [12], while machine-learning approaches have been explored for predicting brain folding [13] and optimizing noninvasive stimulation strategies [14].

Substantial progress has been made in understanding and modeling the material properties of brain tissue over the past decades. Early experimental investigations revealed pronounced regional variation in stiffness [15], motivating finite element models that included basic regional differentiation [6].

Subsequent studies demonstrated strong dependencies of mechanical properties on age and species, in addition to the anatomical region [16]. Detailed mechanical regional characterizations of human tissue have only recently become available [17, 18] and revealed that stiffness varies significantly across regions but exhibits minimal directional dependence, suggesting only a limited mechanical influence of axonal fiber orientation. Experimental studies on nine distinct anatomical regions emphasized the need for region-specific material properties [19]. Corresponding parameters were identified using inverse approaches based on the one-term Ogden model [20] and subsequently applied in simulations of neurosurgical retraction [21]. More recent work extended this concept to a larger number of anatomically distinct regions [22]. Despite their increasing anatomical fidelity, all of these models rely on regionally averaged material parameters derived from ex vivo tissue samples. In parallel, several approaches have emerged that aim to deduce mechanical properties in vivo. A relationship between tissue stiffness and fractional anisotropy (FA) derived from diffusion tensor imaging (DTI) was first reported in [17] and recently confirmed in [23], supporting the use of diffusion MRI as a surrogate for estimating local mechanical properties. DTI has also enabled voxel-wise incorporation of anisotropy in computational models [24], while tractography-based methods further improved the representation of microstructural organization [25]. In addition, magnetic resonance elastography (MRE) has been used to estimate subject-specific, regionally heterogeneous material properties non-invasively, demonstrating the feasibility of personalized biomechanical characterization [26]. Nevertheless, current parameter identification strategies still rely either on indirect imaging correlations or mechanical testing of post-mortem tissue, limiting their applicability in clinical settings. These limitations raise a fundamental question: whether personalized, spatially varying brain material properties can be estimated non-invasively, in vivo, to enable truly patient-specific biomechanical digital twins.

In this study, we explore the feasibility of using diffusion-weighted MRI, and specifically voxel-wise FA values, to infer subject-specific stiffness variations throughout the brain. Leveraging prior work identifying nine mechanically distinct brain regions [17, 20], we determine a linear relationship between regional FA and shear modulus, and apply this mapping to construct a continuous voxel-wise stiffness distribution for the entire human brain, based on the findings of [23]. This approach allows us to estimate local mechanical properties directly from imaging data without invasive or non-standard procedures.

To evaluate the impact of locally varying tissue properties on full-scale human brain simulations, we construct a 3D finite element model of the brain and compare two methods: a region-wise 9-region (9R) parameterization, in which material parameters are constant within each anatomical region, and a voxel-wise FA-based parameterization, in which parameters vary continuously according to local FA values. Both parameterizations are applied to simulate cerebral atrophy. While the goal is not to model atrophy mechanisms in detail, this provides a consistent framework for assessing how different material parameters influence predicted deformation and stress. The comparison serves as a proof of concept for the use of FA-derived stiffness distributions in generating personalized biomechanical parametrization of the human brain.

## 2. Methods

### 2.1. MRI Data Acquisition and Preprocessing

#### Subject and imaging

A total of 15 brains were scanned using a Siemens MAGNETOM Cima.X scanner equipped with a 64-channel head coil. Each subject underwent both a T1-weighted anatomical scan and a diffusion-weighted imaging (DWI) scan, acquired using a 2D Echo-Planar Imaging (EPI) diffusion sequence.

#### T1-weighted scan

The T1-weighted anatomical images were acquired sagittally. The echo time (TE) was 44 ms and the repetition time (TR) between successive excitations was 3500 ms. The data were stored in NIFTI format with a matrix size of 208 × 240 × 256 and an isotropic voxel size of 1 mm × 1 mm × 1 mm, corresponding to a field of view (FOV) of 208 mm × 240 mm.

#### Diffusion-weighted scan

Diffusion-weighted images were acquired with a TR of 3.5 s in 30 diffusion directions, each repeated twice (*p*^2^ acquisition) to enhance signal-to-noise ratio. The diffusion weighting was set to a *b*-value of 1000 s/mm^2^. Images were acquired with a matrix size of 128 × 128, yielding an in-plane resolution of 1.719 mm × 1.719 mm. Slice thickness was 2 mm with an inter-slice spacing of 0.6 mm (effective spacing: 2.6 mm). A total of 54 slices were collected, resulting in a dataset of size 128 × 128 × 54 with voxel dimensions of 1.719 × 1.719 × 2.60 mm^3^. Fractional anisotropy (FA) values were calculated using the manufacturer’s software.

#### Segmentation and Registration

The T1-weighted anatomical images were processed using FreeSurfer image analysis suite^1^’s recon-all automated pipeline, which performs cortical and subcortical segmentation. This procedure yields a 3D label map, in which each voxel is assigned to a specific anatomical region.

The diffusion-derived FA map was subsequently aligned to the T1 anatomical space using FreeSurfer’s `mri_robust_register` tool. This registration step ensures that the FA volume is rigidly aligned and resampled onto the same spatial grid and coordinate system as the T1-weighted MRI.

Following registration, the FA image was masked to include only brain tissue, thereby excluding non-brain structures, such as skull and dura mater.

The final outcome of this preprocessing is a pair of spatially aligned 3D arrays on an identical voxel grid: one containing anatomical region labels (from the T1 segmentation), and the other containing voxel-wise FA values (from the DWI acquisition).

### 2.2. 3D Brain Models

#### Mesh Generation and Coarsening

The segmented T1 volume was converted into a structured hexahedral mesh with an initial resolution of 1 × 1 × 1 mm^3^ elements. To reduce computational cost, this mesh was coarsened to 2 × 2 × 2 mm^3^ elements, while preserving the overall brain geometry. Coarsening was performed using an in-house Python meshing script^2^ that applies a majority-vote strategy: within each 2 × 2 × 2 block, the most frequently occurring region label is assigned to the entire coarse element. A previous mesh analysis demonstrated that simulations performed on both the 1 × 1 × 1 mm^3^ and 2 × 2 × 2 mm^3^ meshes yield comparable results, supporting the validity of this approach [22]. The script also applied Laplacian surface smoothing to all anatomical interfaces, including inter-regional boundaries and the outer brain surface, and extracted region-wise values needed for material assignment. The final mesh contained 248,376 hexahedral elements and 790,320 degrees of freedom. This discretization was used for both region-wise and voxel-wise FA-based parameterization simulations.

Figure 2 shows FA values mapped on the same brain using the fine mesh (left) and the corresponding mesh after coarsening (right). In the coarsening step, rather than computing a simple average, which can create large discrepancies at interfaces where FA values change sharply, the same majority-value strategy as above is used, preserving the spatial patterns and regional trends in FA, and maintaining the characteristic structure of the high and low-FA regions. In addition, a threshold was applied to high FA values, to prevent negative shear moduli values, particularly affecting the corpus callosum, as detailed in Section 2.4.2. This leads to a more homogeneous FA field within some regions, as illustrated by the zoomed-in view of the corpus callosum in Figure 2. Naturally, differences arise between the fine and coarse models. Figure 3 shows boxplots of the FA distributions in the uncoarsened mesh, before thresholding, and in the coarsened mesh, after thresholding. The numbers shown above each boxplot correspond to the number of elements in each region for the fine and coarse meshes. As expected, the coarsening and thresholding steps substantially reduce the number of voxels per region. Despite this reduction, the overall regional FA trends are well preserved, consistent with the spatial comparison in Figure 2. Before thresholding, several regions display wide FA distributions, which naturally reduce once high FA values are clipped. For most regions, the boxplots before and after coarsening are closely aligned, and the medians show good agreement. The main exception is the corpus callosum. This region contains a large proportion of FA values above the threshold, leading to a pronounced difference between the original median (≈ 0.72) and the post-threshold median (≈ 0.59). This behavior aligns with the spatial differences noted earlier and highlights the sensitivity of this region to the thresholding step. These deviations warrant further investigation and should be addressed in future studies.

Consistent with reported subarachnoid cerebrospinal fluid (CSF) thicknesses of approximately 1-5 mm [27] and its increase with aging [28], a 3 mm CSF layer was added to the outer brain surface. The CSF was meshed together with the brain and modeled as an ultrasoft, compressible solid with a stiffness one order of magnitude lower than cortical tissue, following the approach of prior studies [29, 22]. CSF stresses remained negligible relative to those in brain tissue, and no artificial stress concentrations were observed. The external CSF boundary was assigned a homogeneous Dirichlet boundary condition, corresponding to the inner surface of the skull [29].

Motivated by the considerations in [20], we defined nine governing anatomical regions in brain tissue: cortex (C), basal ganglia (BG), brainstem (BS), cerebellum (CB), corpus callosum (CC), corona radiata (CR), hippocampus (Hi), amygdala (Am), and midbrain (M). These regions, shown in Figure 1, form the basis of the region-wise parameterization. The CSF is omitted from Figure 1 for clarity. A figure comprising the mesh and the CSF is included in Figure A.13 in Appendix Appendix A.

**Figure 1:**
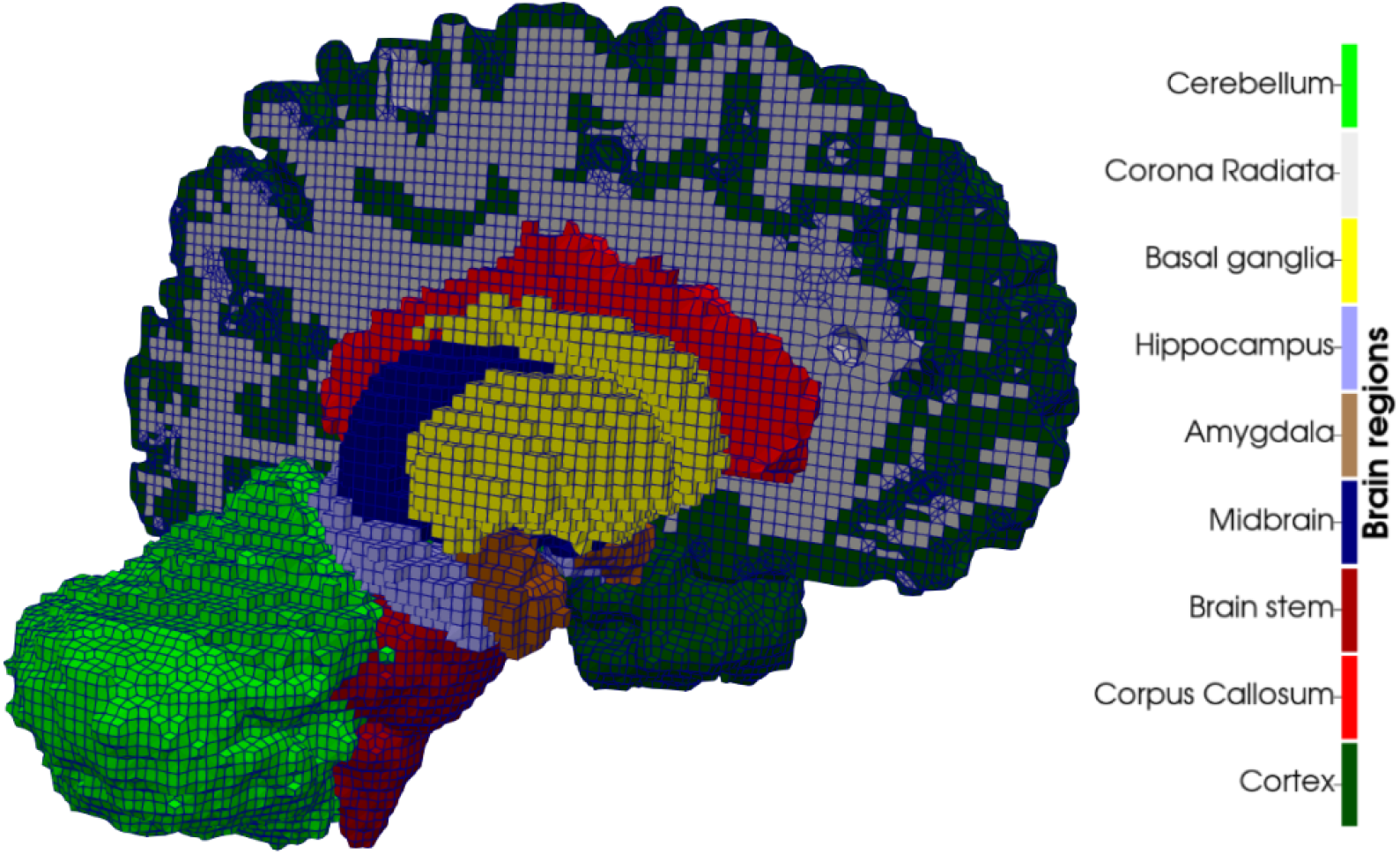
Regional segmentation for the region-wise (9R) parameterization. The surrounding cerebrospinal fluid (CSF) layer is omitted for clarity.

**Figure 2:**
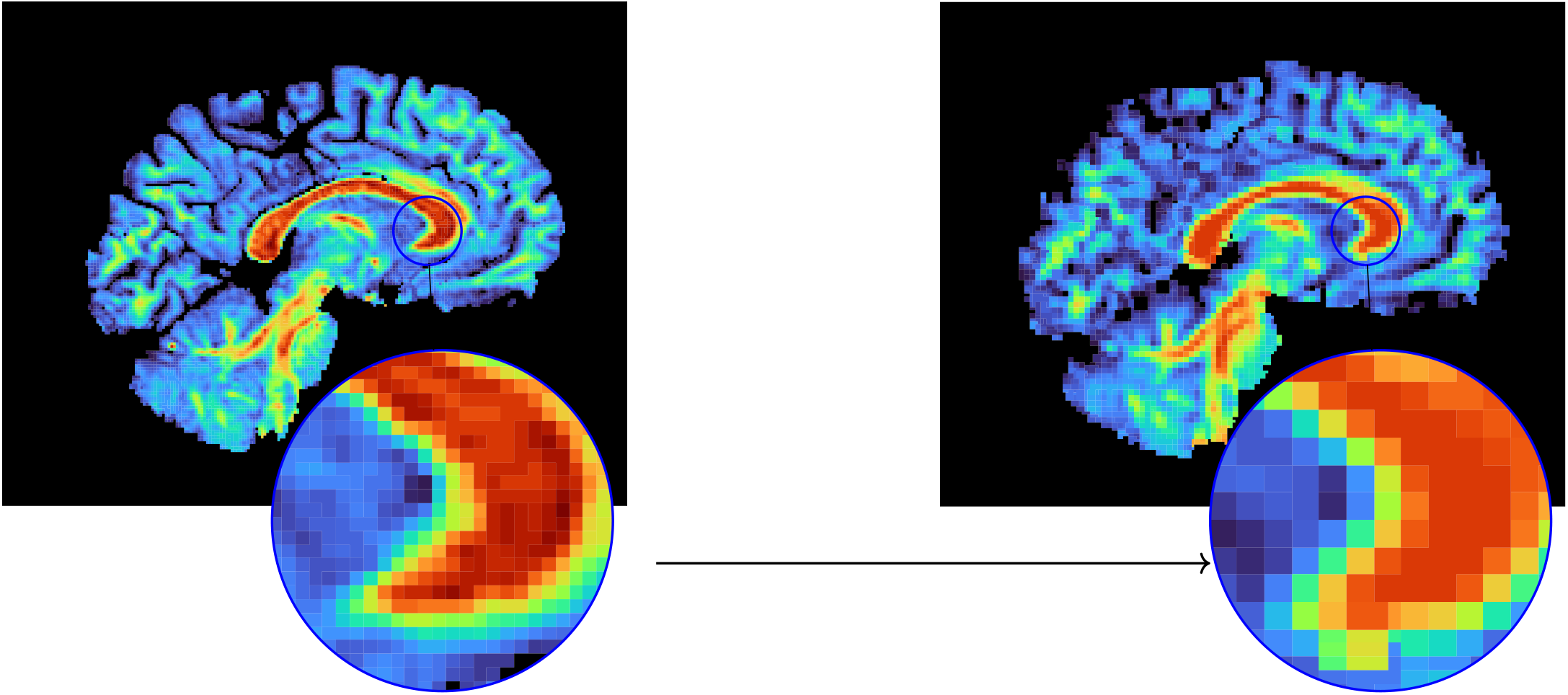
Fractional Anisotropy (FA) map before (left) and after (right) coarsening. Each 2 × 2 × 2 voxel block from the original 256^3^ image is merged into a single voxel in the 128^3^ coarsened volume. Zoom on Corpus callosum shows a more homogeneous FA field within this region for the coarse mesh.

**Figure 3:**
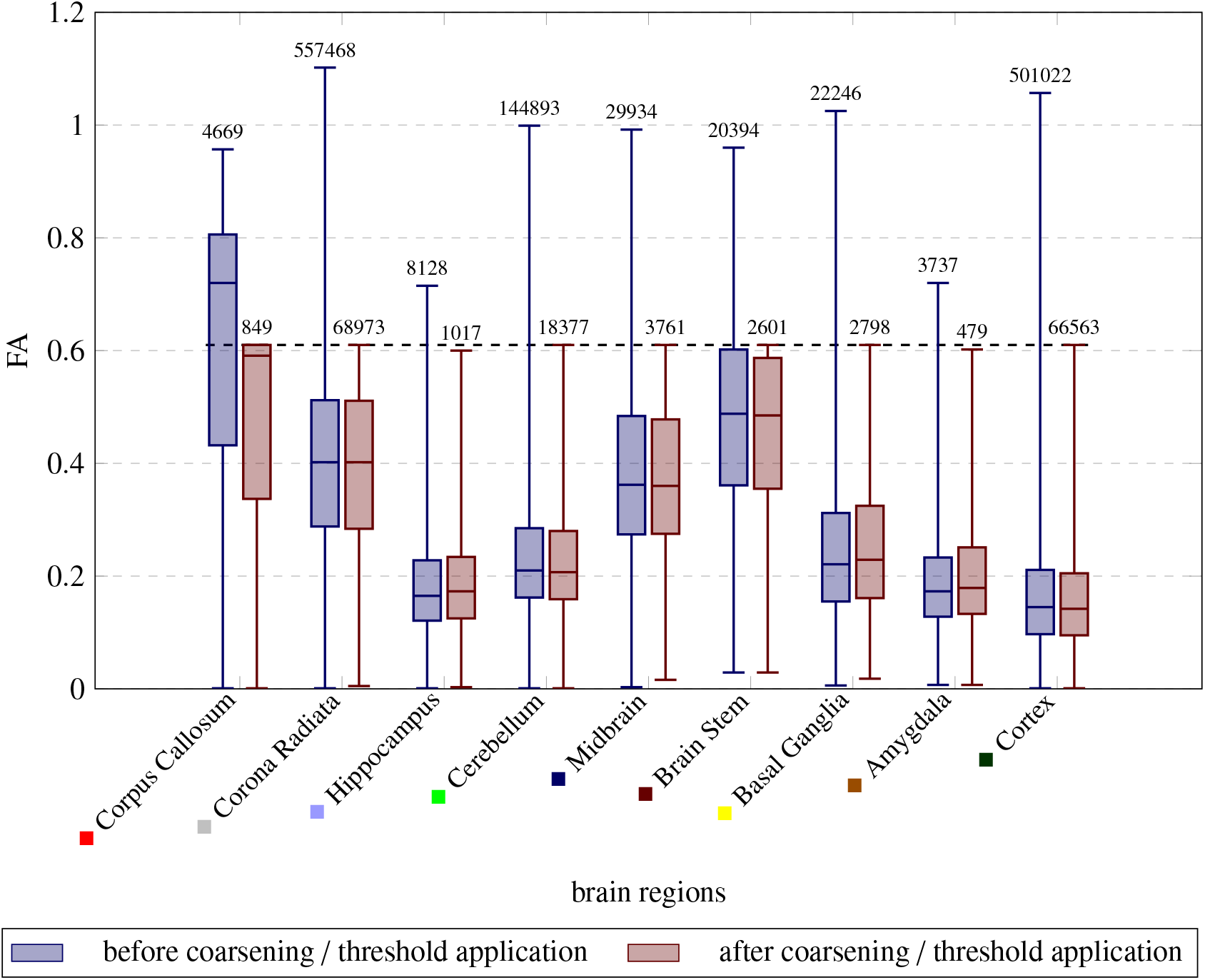
Blue boxplots show the voxel-wise FA distributions in the refined (pre-coarsening) mesh, and red boxplots show the corresponding distributions after coarsening and thresholding. The boxes represent the interquartile range (25th–75th percentiles), and the whiskers span the full range of the data. The numbers above each boxplot indicate the number of elements in that region before and after coarsening and thresholding. The dashed horizontal line at FA = 0.61 marks the applied threshold.

### 2.3. Constitutive Model

Brain tissue is modeled as a compressible hyperelastic solid using a modified one-term Ogden strain-energy formulation, which has been shown to accurately capture the nonlinear, compression-tension asymmetric and rate-independent mechanical response of brain tissue under a wide range of loading conditions [17]. Viscous effects are neglected, consistent with prior studies focusing on quasi-static or rate-independent loading regimes.

We model brain tissue as mechanically isotropic. Although magnetic resonance elastography (MRE) studies have reported anisotropic mechanical behavior in white matter [30], and recent large-strain experiments have revealed anisotropy in certain structures, most prominently in the corpus callosum [31], direction-dependent effects remain much less pronounced at the whole-brain scale and for the strain levels considered here. Furthermore, no region-wise anisotropic material parameters are currently available for the human brain. For these reasons, an isotropic hyperelastic model is adopted.

Following [32], we decompose the total strain-energy density into isochoric and volumetric contributions,

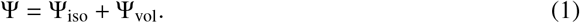

Let ***F*** denote the deformation gradient and *J* = det ***F*** its Jacobian. Using the isochoric principal stretches 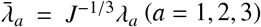, the isochoric part is given by

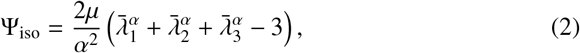

where *µ* is the shear modulus and *α* represents the degree of nonlinearity and compression-tension asymmetry.

The volumetric part is defined as

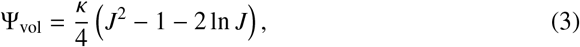

where the bulk modulus κ is related to the shear modulus *µ* and Poisson’s ratio *ν* via

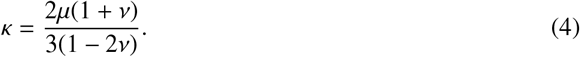

The Cauchy stress tensor follows from finite hyperelasticity as

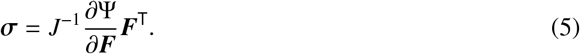

### 2.4. Material Parameters

All simulations were performed on the same brain model generated as described above. Two distinct strategies for assigning material properties were considered: a region-wise parameterization and a voxel-wise FA-based parameterization, as detailed in the following.

#### 2.4.1. Region-wise parameterization

Each of the nine anatomical regions in the 9R parameterization (see Figure 1) was assigned a set of homogeneous material parameters, a shear modulus *µ* and a nonlinearity parameter *α*, summarized in Table 1. These values were obtained by fitting a modified Ogden hyperelastic model to experimental stress-strain data from human brain tissue samples extracted from each region in [20]. Cerebrospinal fluid (CSF) and ventricular regions were assigned very low stiffness values to emulate their fluid-like behavior. The choice of the 9R parameterization follows a sequence of earlier studies that evaluated how increasing levels of tissue heterogeneity affect brain mechanics simulations [21, 33, 22].

**Table 1:** Material parameters averaged over anatomical regions, extracted from [20].

| Gov. region | Preconditioned<br>$\nu = 0.45$ ● | | Preconditioned<br>$\nu = 0.49$ ● | | Unconditioned<br>$\nu = 0.45$ ● | | Unconditioned<br>$\nu = 0.49$ ● | |
| --- | --- | --- | --- | --- | --- | --- | --- | --- |
| | $\alpha$ | $\mu$ | $\alpha$ | $\mu$ | $\alpha$ | $\mu$ | $\alpha$ | $\mu$ |
| Amygdala | -19.67 | 257.05 | -14.16 | 193.34 | -14.74 | 386.97 | -10.10 | 293.06 |
| Basal ganglia | -18.95 | 252.15 | -15.05 | 169.67 | -15.90 | 338.67 | -11.49 | 246.21 |
| Brain stem | -20.41 | 233.12 | -19.59 | 118.06 | -16.82 | 336.44 | -13.60 | 225.08 |
| Cortex | -19.27 | 281.75 | -16.72 | 175.21 | -16.16 | 392.24 | -12.16 | 285.97 |
| Cerebellum | -23.22 | 177.49 | -18.79 | 115.36 | -19.49 | 255.38 | -14.08 | 196.55 |
| Corpus callosum | -14.94 | 61.66 | -13.72 | 37.64 | -12.82 | 95.43 | -9.11 | 67.82 |
| Corona radiata | -19.70 | 157.09 | -19.08 | 87.58 | -16.93 | 218.75 | -13.55 | 146.54 |
| Hippocampus | -17.37 | 184.21 | -13.09 | 129.96 | -13.25 | 292.58 | -9.23 | 218.74 |
| Midbrain | -18.68 | 215.70 | -15.15 | 137.67 | -15.11 | 314.92 | -10.89 | 227.69 |

#### 2.4.2. Voxel-wise FA-based parameterization

*Linear fit between FA and shear modulus*. For each of the nine governing regions, the average fractional anisotropy (FA) was computed across the cohort of 15 brains. These regional mean FA values were paired with the corresponding regional shear modulus values *µ* from the 9R dataset, in order to identify a mapping from diffusion-derived FA to mechanical stiffness.

A linear regression was performed to fit the region-wise shear modulus as a function of the region-averaged FA,

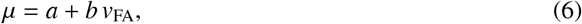

where *µ* is the shear modulus (piecewise constant per region in the original dataset), and *v*_FA_ is the region-wise averaged value of the FA.

The Pearson correlation coefficient was found to be high (|*r*| > 0.8) and statistically significant (*p* ≤ 0.002), demonstrating a strong negative correlation between FA and *µ*. Regions with high FA values exhibited lower shear moduli, whereas regions with low FA values exhibited higher moduli. No statistically significant correlation was observed for the nonlinearity parameter *α*. A detailed analysis of these relationships is provided in Section 3.1.

#### Local material properties

Motivated by the strong negative correlation between the FA value and the shear modulus, we developed a voxel-wise FA-based human brain parameterization, where material parameters vary continuously throughout the brain. Each finite element is assigned its own stiffness value by evaluating the linear FA-*µ* mapping introduced in Equation 6 based on the element’s local FA value. This yields a spatially heterogeneous shear modulus field that reflects microstructural differences captured in the DWI.

Since high FA values could produce negative shear moduli under the linear mapping, an upper FA threshold was imposed to ensure physically meaningful stiffness values. The nonlinearity parameter *α* is kept uniform (piecewise constant per region), consistent with the region-wise parameterization, due to the absence of a significant FA-*α* correlation.

As in the region-wise parameterization, CSF and ventricular regions were assigned significantly lower stiffness values to represent fluid-like mechanical behavior.

### 2.5. Modeling Cerebral Atrophy

To model cerebral atrophy, we closely follow the approach introduced in [22]. The method is implemented in deal.II, and the corresponding code is publicly available^3^. Atrophy is introduced kinematically through a multiplicative decomposition of the deformation gradient into elastic and atrophy parts [34, 29],

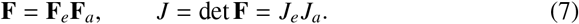

We assume isotropic shrinkage of the form

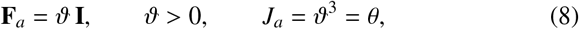

so that the elastic response is evaluated from 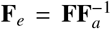. The atrophy-weighted strain-energy density becomes

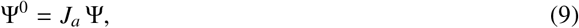

such that stresses depend solely on the elastic quantities **F**_*e*_ and *J*_*e*_.

Following [29], the volume-change ratio θ evolves according to

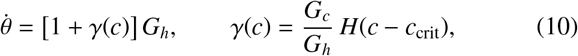

i.e.,

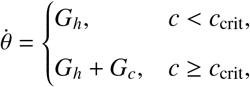

where *G*_*h*_ and *G*_*c*_ denote region-dependent healthy and pathological atrophy rates for gray matter, white matter, and the hippocampus. The atrophy parameters used in this study are based on Alzheimer-type progression [35] and are summarized in Table 2. Values are reported as percentages per year, and negative values correspond to shrinkage.

**Table 2:** Healthy (*G*_*h*_) and pathological (*G*_*c*_) atrophy rates for grey matter, white matter, and hippocampus, consistent with Alzheimer-type progression.

| | $G_h$ [%/year] | $G_c$ [%/year] |
| --- | --- | --- |
| **Grey matter** | -0.10 | -0.20 |
| **Hippocampus** | -0.10 | -0.20 |
| **White matter** | -0.15 | -0.35 |

White matter atrophy is assumed to exceed gray matter atrophy, in agreement with earlier findings [36, 37]. A synthetic protein concentration field *c*(**x**) was prescribed, peaking in the hippocampal region and decaying toward the cortex. The threshold *c*_crit_ leads to accelerated atrophy in the hippocampus and adjacent temporal regions.

To model progressive shrinkage, the problem is solved incrementally in a quasi-static framework. At each time step *n* → *n*+1, the atrophy measure θ ^*n*+1^ is updated and mapped into the atrophy tensor

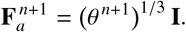

The elastic equilibrium configuration is then computed from **F** = **F**_*e*_**F**_*a*_ with the updated elastic deformation gradient 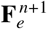. Because physiological atrophy occurs over months to years, viscous and porous effects are neglected and each increment is treated as a quasi-static elastic equilibrium.

### 2.6. Postprocessing and Statistical Analysis

#### Postprocessing

Both region-wise and voxel-wise simulations were performed on the same subject-specific brain mesh. The simulation outputs included displacement fields, principal stretches, maximum shear strain, hydrostatic stress, and von Mises stress.

For analysis, all quantities were aggregated over the nine anatomical regions defined in the 9R parameterization model. For each output variable, we computed the region-wise differences between the region-wise and voxel-wise simulations to quantify how the material parameters affected the predicted mechanical response.

#### Statistical Analysis

Because both simulations were conducted on the same mesh, the resulting element-wise data are inherently paired. For each region, we computed the element-wise differences

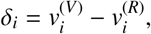

where 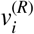 denotes the value from the region-wise simulation and 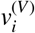 the corresponding value from the voxel-wise simulation for element *i*.

The distributions of stress and stretch measures within regions deviated from normality (see Figures B.14, B.15, B.16 and B.17 in Appendix Appendix B). Therefore, non-parametric statistical tests were employed. Specifically, the Wilcoxon signed-rank test was used to assess whether the differences between the two parameterizations were statistically significant. The test consistently yielded *p* < 0.001, indicating differences in the distributions.

To complement *p*-values and quantify the magnitude of differences, we calculated Cohen’s *d*, defined as:

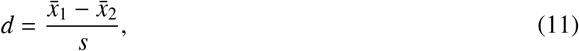

where 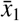 and 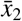 are the sample means of the two parameterizations and *s* is the pooled standard deviation. All statistical analyzes were performed using the statistical functions module (scipy.stats, version ≥ 1.7.0) in Python.

## 3. Results and Discussion

### 3.1. FA-based stiffness prediction

Linear regressions were performed as described in Section 2.4.2 for four different material configurations: preconditioned tissue response with a Poisson’s ratio *ν* = 0.45, preconditioned tissue response with a Poisson ratio *ν* = 0.49, unconditioned tissue response with a Poisson ratio *ν* = 0.45, and unconditioned tissue response with a Poisson ratio *ν* = 0.49. Table 3 summarizes the linear relationships identified between the shear modulus *µ* and the FA values for the four datasets. For each case, a linear model was fitted, and the corresponding coefficient of determination *R*^2^ and Pearson correlation *p*_*r*_ value are reported. Figure 4, left, illustrates the correlation exemplary for the pre-conditioned dataset with *ν* = 0.45, where a clear negative linear relationship is visible between the regional FA values and the shear modulus *µ*, with regions of higher stiffness exhibiting lower FA. Across all datasets, we observe a statistically significant correlation between *µ* and FA, with *R*^2^ values ranging from 0.642 to 0.753 and *p*_*r*_ values between 0.006 and 0.009. The strongest linear association is found for the preconditioned dataset with *ν* = 0.49 (*R*^2^ = 0.753), indicating that preconditioning combined with a higher Poisson’s ratio improves the predictability of *µ* from the FA value. In contrast, the pre-conditioned dataset with *ν* = 0.45 shows the lowest correlation (*R*^2^ = 0.642), although the relationship remains significant.

**Table 3:** Overview of the different linear fit functions for the four *µ* value datasets and the corresponding statistics of the Pearson test.

| Dataset | Linear mapping | $R^2$ | $p_r$ | Exemplary Application |
| --- | --- | --- | --- | --- |
| Preconditioned, $\nu = 0.45$ | $\nu_{FA} = -0.0017\mu + 0.6236$ | 0.642 | 0.009 | Slow processes such as aging/disease progression; slightly compressible |
| Preconditioned, $\nu = 0.49$ | $\nu_{FA} = -0.0031\mu + 0.6066$ | 0.753 | 0.002 | Slow processes such as aging/disease progression; quasi incompressible |
| Unconditioned, $\nu = 0.45$ | $\nu_{FA} = -0.0012\mu + 0.6470$ | 0.669 | 0.007 | Faster processes such as impact loading; slightly compressible |
| Unconditioned, $\nu = 0.49$ | $\nu_{FA} = -0.0016\mu + 0.6465$ | 0.683 | 0.006 | Faster processes such as impact loading; quasi-incompressible |

**Figure 4:**
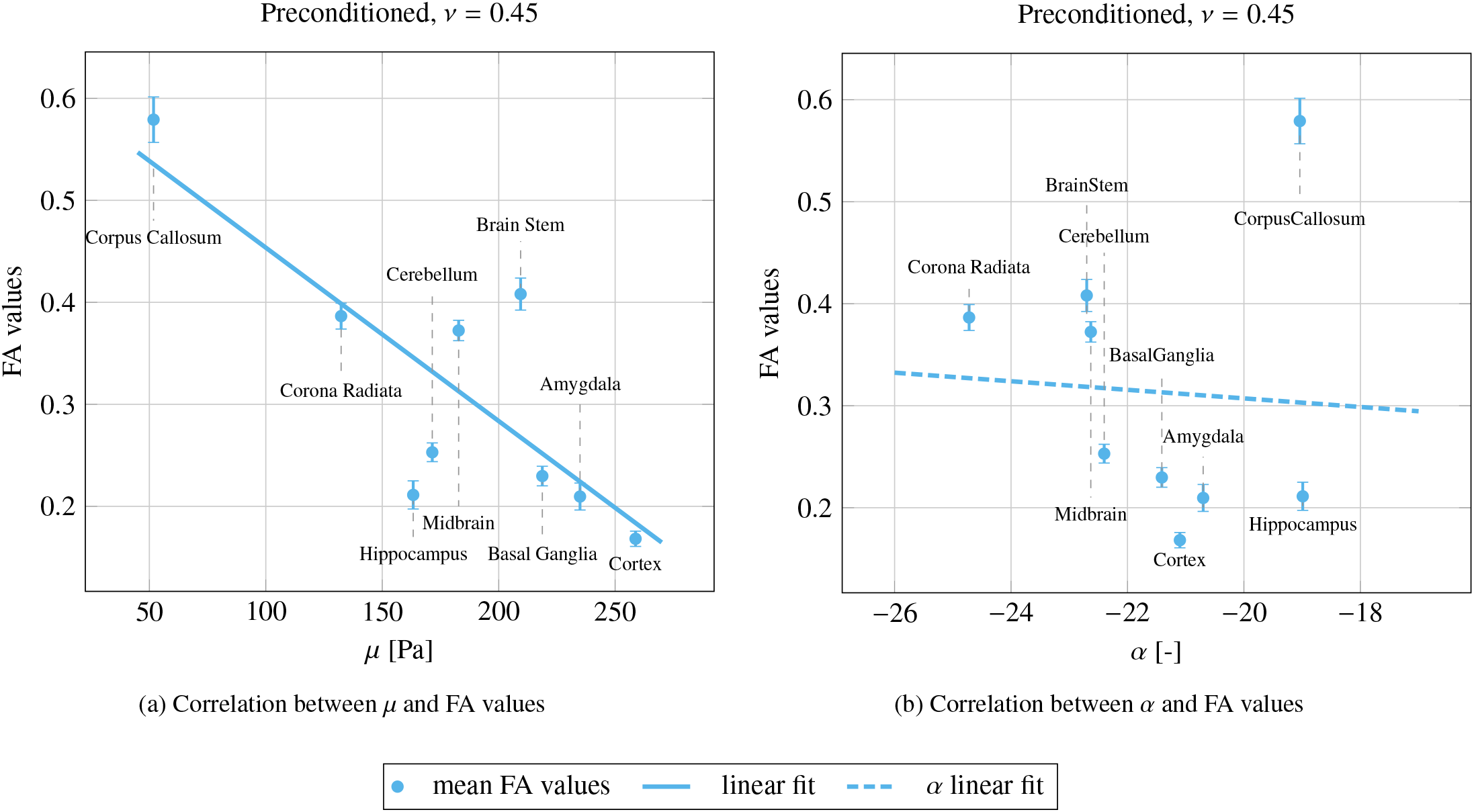
Comparison between *µ* (left) and *α* (right) correlations for the preconditioned, *ν* = 0.45 dataset.

Across the individual anatomical regions, the cortex, basal ganglia, and amygdala align best with the linear negative correlation. Larger deviations occur for the brain stem, hippocampus and cerebellum, which may reflect local structural heterogeneity, not raised by the mechanical testing.

The appropriate choice of one of the four parameter sets depends on the loading conditions relevant for the desired application. The right column of Table 3 summarizes relevant biomechanical scenarios for each parameter set. Preconditioned mappings are more representative of slow, quasi-static processes such as tissue aging or disease progression, while unconditioned mappings better reflect rapid mechanical loading conditions, such as impact or trauma. For all subsequent simulations, we adopt the *preconditioned* material parameters with a Poisson ratio of *ν* = 0.45. This choice represents a slightly compressible tissue response, which is appropriate for modeling physiological volume loss during progressive atrophy. Nonetheless, alternative values of *ν* may be considered for applications involving different loading regimes.

Table C.4 in Appendix Appendix C presents the corresponding linear regressions between the region-wise nonlinearity parameter *α* and the regional FA values. Across all conditions, the correlation coefficients are close to zero, indicating no meaningful relationship between *α* and FA values. Accordingly, Figure 4, right, shows no discernible relationship between FA and the nonlinearity parameter *α*. These results confirm that FA is predominantly sensitive to regional variations in *µ* rather than *α*. Hence, *α* is not derived from FA in the voxel-wise FA-based parameterization. Instead, we retain the region-wise values obtained from [20]. This ensures spatial heterogeneity in *α*, which has shown to produce more realistic simulations [21].

### 3.2. Region-wise versus voxel-wise FA-based shear moduli in brain atrophy simulations

Figure 5 compares the regional segmentation of the brain and the corresponding material parameter distributions for both the region-wise and FA-derived voxel-wise parameterizations. Panel (a) shows the nine anatomical regions used in the 9R parameterization, which serve as the basis for assigning homogeneous material parameters within those regions. Panel (b) presents the resulting shear modulus *µ* distribution for the region-wise parameterization. As expected, each region appears uniform, reflecting the homogeneous parameter assignment. Distinct stiffness contrasts emerge between regions, with higher *µ* values in structures such as the cortex, basal ganglia, and brain stem, and lower values in white matter regions such as the corpus callosum and corona radiata. This parameterization captures macroscopic differences in tissue but does not account for intraregional variability.

**Figure 5:**
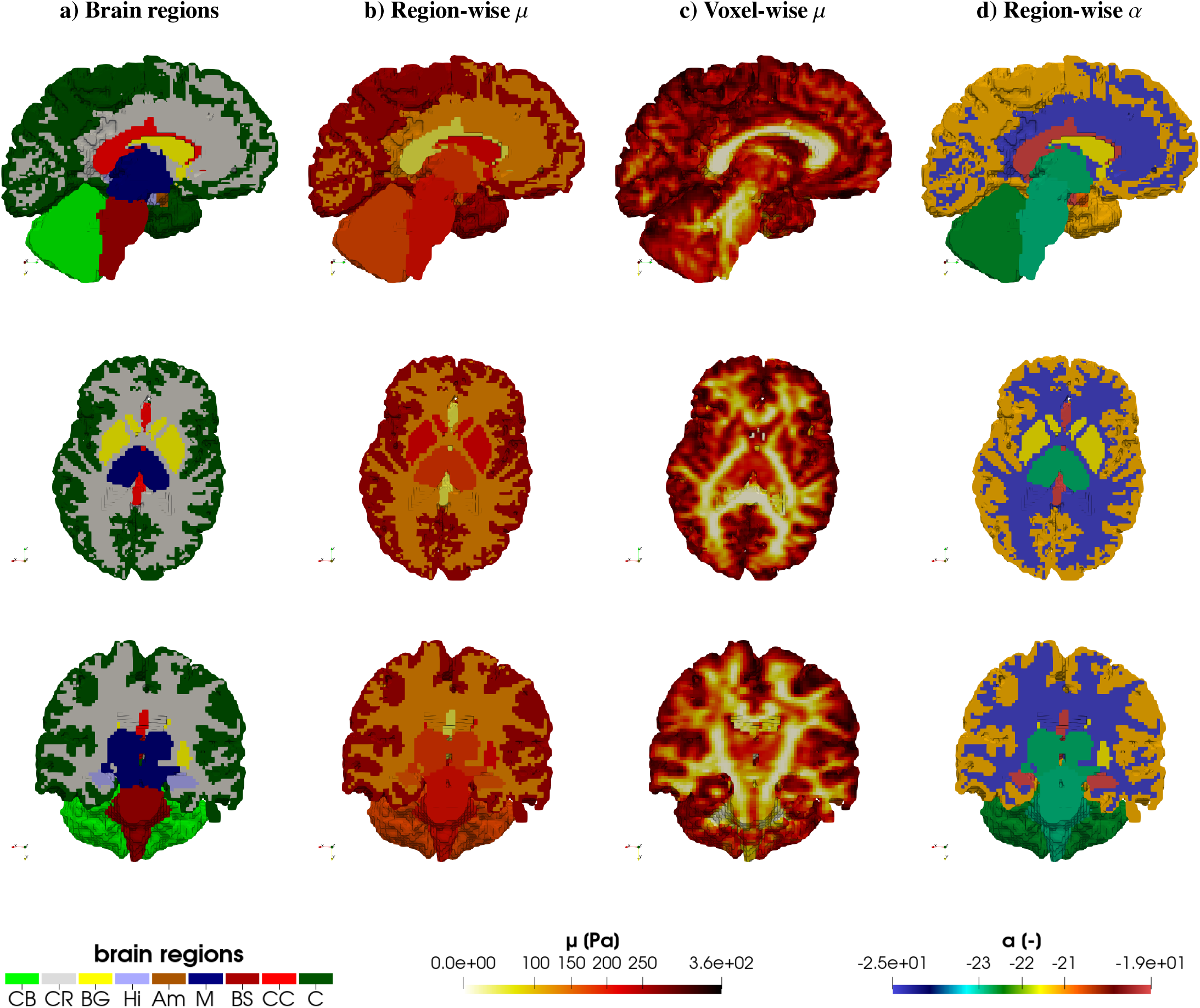
Sagital, transverse and coronal slices of the brain showing a) anatomical regions: CB–cerebellum, BG–basal ganglia, Hi–hippocampus, Am–amygdala, M–midbrain, BS–brain stem, CC–corpus callosum, C–cortex, CR–corona radiata, b) the region-wise parameterization shear modulus *µ* distribution, c) the FA-derived voxel-wise parameterization shear modulus *µ* distribution, and d) the non linearity parameter *α* distribution used in both parameterizations.

In contrast, panel (c) illustrates the distribution of the voxel-wise shear modulus *µ* derived from the FA–*µ* linear mapping. This produces a heterogeneous spatial pattern, with obvious variations within each region. Cortical regions appear stiffer, with *µ* values reaching 360 Pa, while the corona radiata, brain stem and midbrain appear softer. Compared to the region-wise map, the FA-wise distribution reveals anatomical detail along white matter tracts and local gradients within individual structures. While finer-scale heterogeneity is captured, the overall regional stiffness trends remain consistent with those observed in the 9R parameterization. This highlights the ability of FA-derived material parameters to capture microstructural heterogeneity, allowing the identification of localized soft or stiff areas that are not accounted for in the region-wise parameterization.

Panel (d) displays the region-wise nonlinearity parameter *α*, which is spatially homogeneous within each anatomical region. As no meaningful correlation was found between FA and *α*, this parameter remains region-wise for both parameterizations. As a result, the mechanical nonlinearities in the voxel-wise FA-based parameterization inherit the same coarse regional structure as the region-wise parameterization, while only the shear modulus varies at the voxel scale.

Figure 6 illustrates the range of shear modulus values, *µ*, for each brain region obtained (1 - filled points) from fitting the response of individual experimentally tested specimens separately in [20] and (2 - boxplots) from the region-wise shear modulus values obtained from fitting the average response of all experimentally tested specimens from one region in [20] (values used for the region-wise simulations) along with the voxel-wise FA-based parameterization introduced in Section 2.4.2.

**Figure 6:**
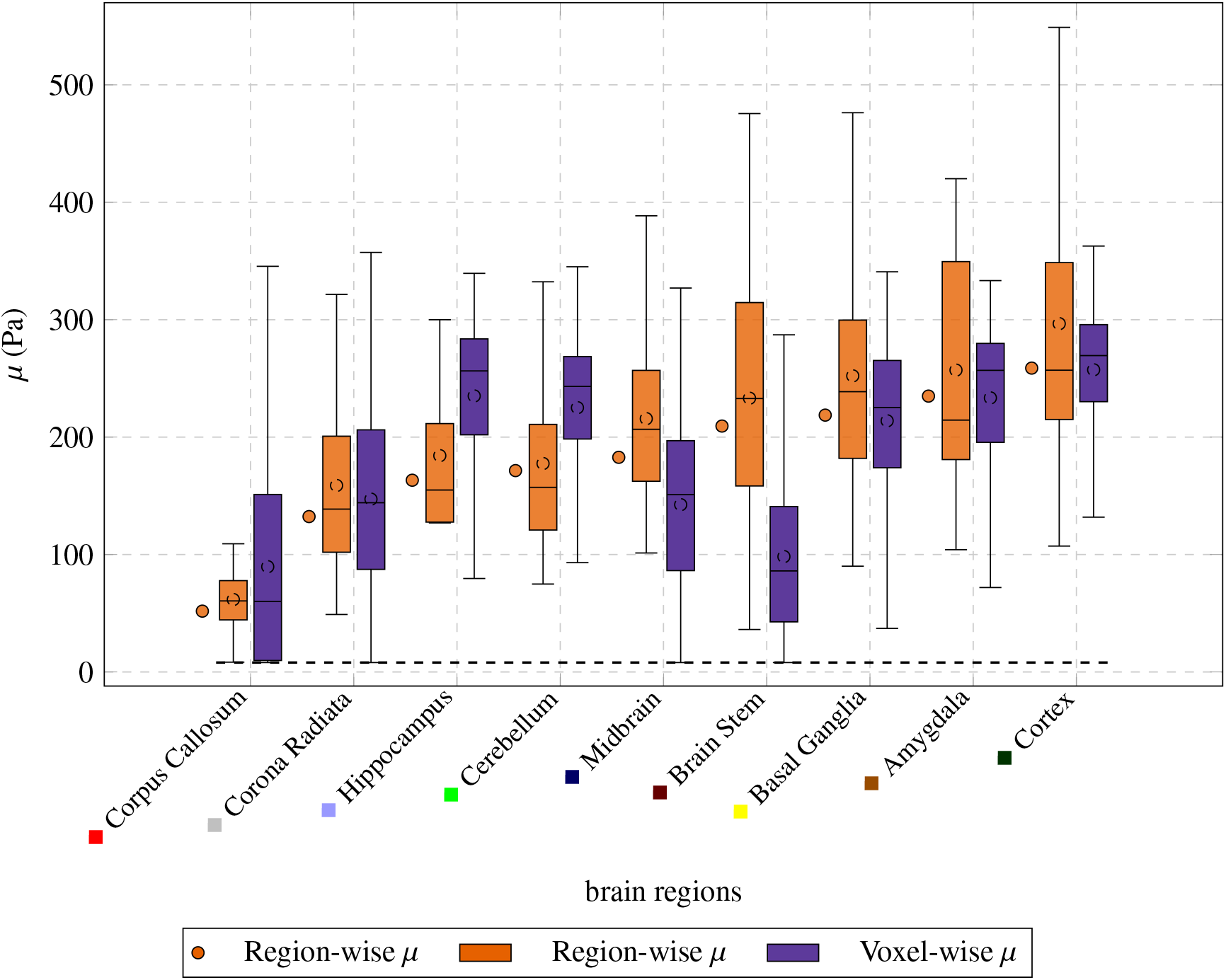
The distribution of *µ* values across all brain regions. Filled points indicate the region-wise parameters used in the simulations, obtained by averaging mechanical responses from multiple samples per region and fitting the material parameter *µ* to the aggregated behaviour. The region-wise boxplots show the variability of *µ* obtained when fitting *µ* independently to each individual sample, as reported in [20]. The voxel-wise boxplots represent the voxel-wise *µ* distributions derived by applying the linear *µ*–FA relationship to each voxel. The dashed horizontal line at *µ* = 8 Pa denotes the lower threshold imposed on FA-derived values to prevent non-physical stiffness.

A dashed horizontal line at *µ* = 8 Pa marks the lower threshold imposed to prevent negative stiffness values in the FA-based dataset, as described in Section 2.4.2. The plot reveals substantial variability in *µ* both across and within brain regions.

The region-wise parameters (filled points) lie within the interquartile ranges of the region-wise boxplots and are close to the mean shear modulus values (used in the region-wise simulations). However, the specimen-based boxplots reveal considerable inter-specimen variability in regions such as the brain stem, amygdala, and cortex, indicating pronounced mechanical heterogeneity that is not captured when a single averaged parameter is assigned per region. In contrast, regions such as the corpus callosum exhibit narrower specimen-based distributions, suggesting more uniform mechanical properties.

The voxel-wise FA-based distributions show distinct trends. Regions including the corpus callosum, corona radiata, midbrain, and brain stem display broad voxel-wise spreads, consistent with the spatial heterogeneity illustrated in Figure 5. By comparison, the FA-based distributions for the cortex and cerebellum are relatively narrow, indicating a more homogeneous FA-derived stiffness distribution within these regions.

The agreement between the experimentally derived and FA-derived distributions is more region-specific. Several regions align well, including the corpus callosum, corona radiata, basal ganglia, amygdala, and cortex. In the hippocampus and cerebellum, the region-wise values lie below the FA-based ranges, implying that these structures are represented as softer in the region-wise parameterization. In the brain stem and midbrain, we observe the opposite effect: the region-wise value is higher than the voxel-wise average, suggesting a stiffer representation in the region-wise parameterization. These differences illustrate that experimentally measured mechanical variability does not always correlate with microstructural variability inferred from FA, underscoring the challenges of assigning material parameters to anatomically complex tissue.

Differences also emerge when comparing medians. The voxel-wise FA-based parameterization yields higher median *µ* values for the hippocampus, cerebellum, amygdala, and cortex, whereas the region-wise parameterization produces higher medians for the midbrain, brain stem, and basal ganglia. These discrepancies indicate that the two approaches can lead to meaningfully different mechanical representations of specific brain regions.

Finally, it is worth noting that the lower stiffness threshold applied to FA values primarily affects the corpus callosum, where high FA values would otherwise map to unrealistically low *µ*. Although the threshold prevents negative values, the resulting *µ* distribution in the corpus callosum remains substantially wider than that obtained from the data experimentally derived in [20].

However, we observe a good agreement in the medians of both distributions.

### 3.3. Predicted volume changes

When simulating the progression of atrophy, both the region-wise and voxel-wise FA-based parameterizations result in similar global brain shrinkage. The FA-based parameterization estimates a total volume reduction of 23%, closely matching the 23.6% shrinkage predicted by the region-wise parameterization, indicating that despite their different parameter approaches, both parameterizations capture the overall magnitude of tissue loss in a similar fashion.

Figure 7 summarizes the percentage volume change at the end of the atrophy simulation for all brain regions (left) and for the ventricles and CSF (right). Among the nine anatomical regions, the largest shrinkage (–33% to –30%) occurs in the internal brain structures, specifically the brain stem, midbrain, amygdala, and basal ganglia, as well as in the cerebellum. This pattern is consistent with the higher protein concentrations prescribed in our model for these regions, and also aligns with clinical observations in Alzheimer’s disease, where these structures are known to undergo substantial degeneration [38, 39]. The cortex exhibits a more moderate reduction in volume (–17%). Overall, the regional shrinkage patterns predicted by the two parameterizations are in good agreement. However, shrinkage is consistently slightly larger in the region-wise parameterization than in the FA-based one, with the largest differences occurring in the brain stem (–33% vs –29.5%) and in the mid-brain (–32.9% vs –30.6%). These discrepancies reflect the differing stiffness profiles imposed by the two parameterizations, observed in Figures 5 and 6.

**Figure 7:**
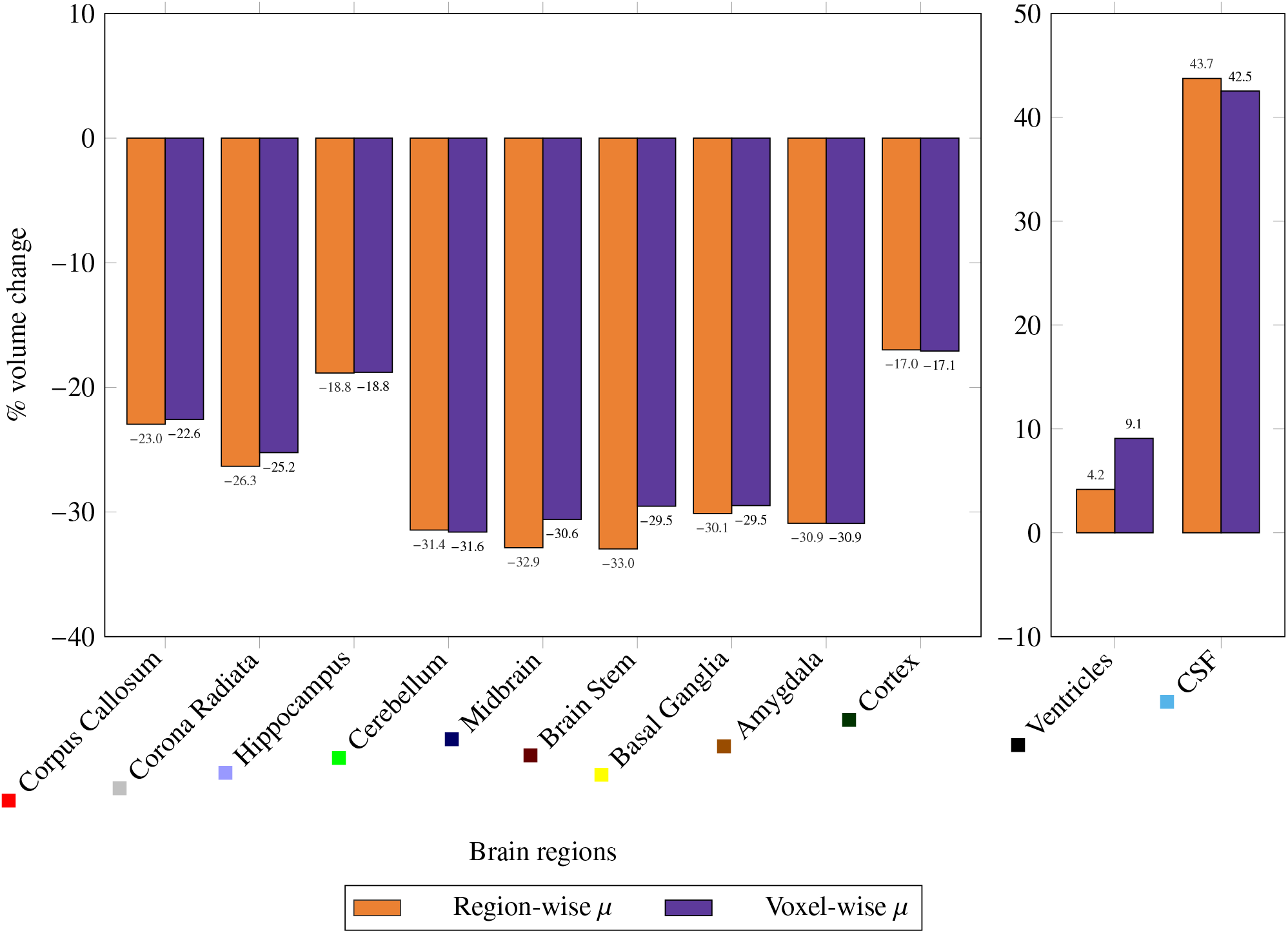
Percentage change in simulated brain region volumes at the final step of the atrophy model, comparing the region-wise (9R) and voxel-wise (FA) parameterizations. Left: 9 brain regions. Right: ventricles and CSF.

A more pronounced divergence between the parameterizations is observed in the predicted ventricular enlargement. Ventricular expansion, a hallmark of neurodegenerative atrophy [35], is +4.2% in the 9R region-wise parameterization but increases to +9.1% in the voxel-wise FA-based parameterization, corresponding to more than twice the enlargement. This difference can be directly attributed to the mechanical stiffness distributions shown in Figure 8, which provides a zoomed-in view of the stiffness values surrounding the ventricles for both parameterizations. In the 9R region-wise parameterization, periventricular regions are assigned piecewise constant stiffness values between approximately 50 and 260 Pa. In contrast, the voxel-wise FA-based parameterization reveals extensive zones of markedly low stiffness (≺ 50 Pa) around the ventricles. These softer regions deform more easily, thereby allowing the ventricles to expand more during the atrophy process. Additionally, the voxel-wise FA-based stiffness distribution exhibits detailed heterogeneity, with localized periventricular areas ranging from 100 to 360 Pa. Such complexity gives rise to regionally varying deformation patterns that cannot be reproduced by the region-wise constant 9R parameterization.

**Figure 8:**
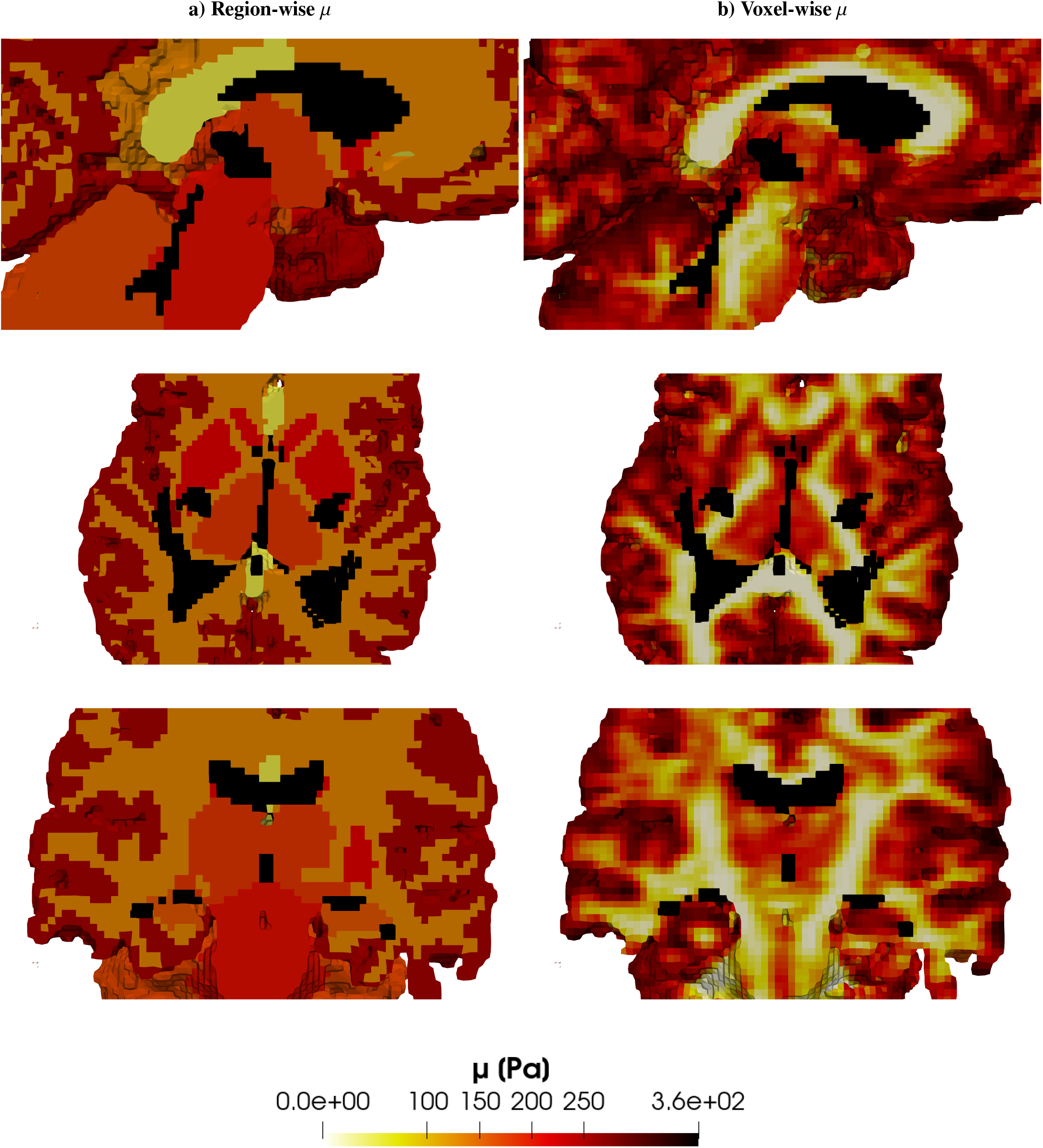
Shear modulus distributions for the (a) region-wise parameterization and the (b) voxel-wise FA-based parameterization, for three anatomical sections of the brain, cropped around the ventricles for better visualization. The ventricles are overlaid in solid black.

Finally, CSF volume increases in both parameterizations as a combined consequence of tissue shrinkage and ventricular expansion. The magnitude of CSF enlargement is similar across the two parameterizations, with a +43.7% increase in the region-wise parameterization and +42.5% in the FA-based parameterization.

### 3.4. Predicted displacements and strains

To assess and compare the mechanical simulation outputs, i.e., displacements, first and third principal stretches as well as the maximum shear, we conducted a comprehensive statistical analysis, as described in Section 2.6.

The results are illustrated using boxplots across the nine anatomical regions.

#### . Displacement

Figure 9 compares the distributions of displacement magnitudes predicted by the region-wise parameterization and the voxel-wise FA-based parameterization for all nine brain regions. Displacement is used as a comparison metric because atrophy is modeled as a prescribed volumetric shrinkage, making displacement a direct measure of the resulting tissue relocation and deformation. This choice is consistent with prior computational studies of brain atrophy, where displacement fields are analyzed to characterize shrinkage-related morphological changes [29, 40]. From a clinical perspective, displacement provides an interpretable measure that directly reflects the macroscopic anatomical changes observed in neuroimaging, such as ventricular enlargement, cortical thinning, and regional tissue displacement.

**Figure 9:**
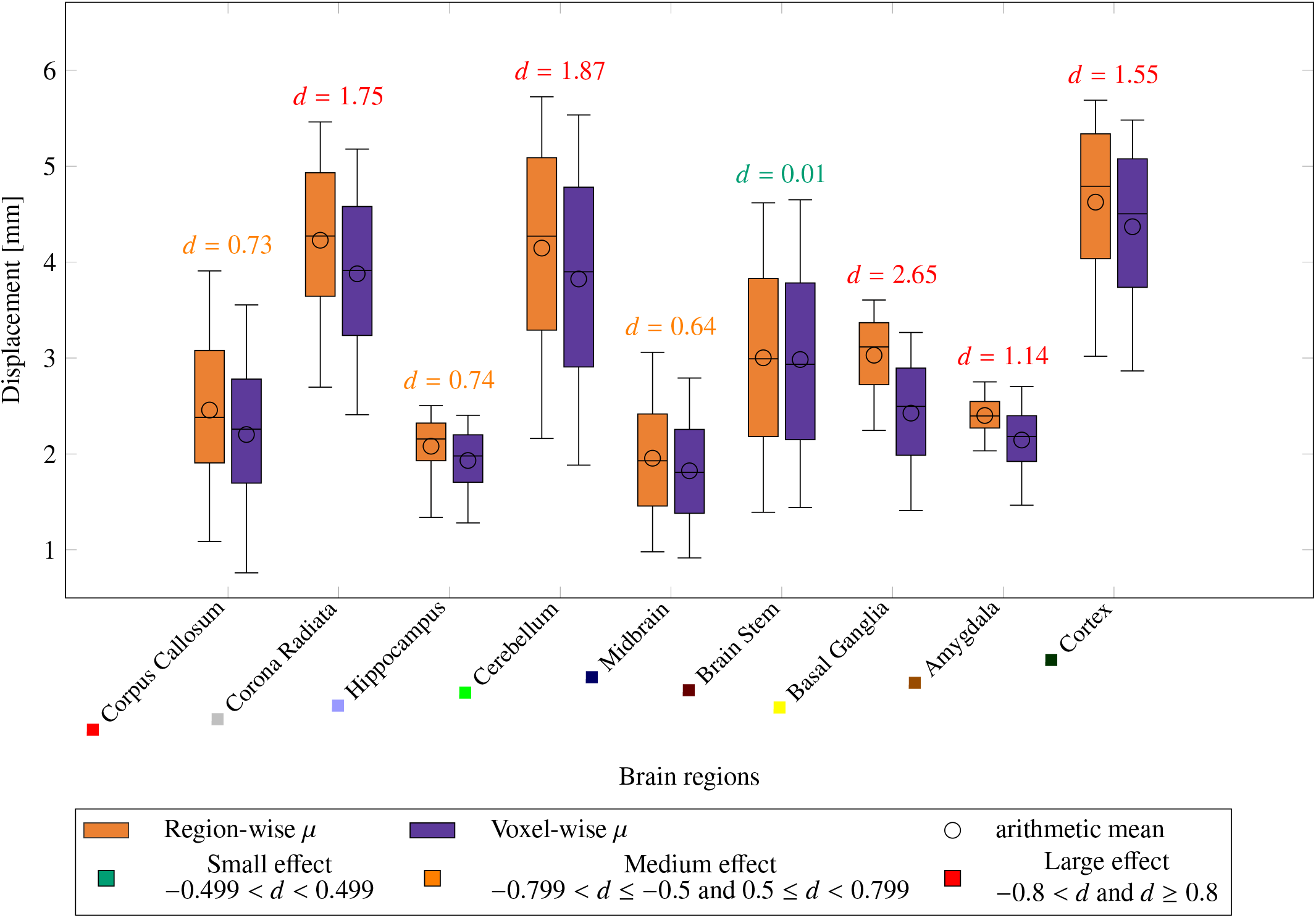
Boxplots illustrate the distributions of the simulated displacement field [mm] across the nine anatomical brain regions for both the region-wise and voxel-wise parameterizations. Whiskers extend to the 5^th^ and 95^th^ percentiles, while the boxes and the central line indicate the 25^th^, 50^th^ (median), and 75^th^ percentiles; the arithmetic mean is indicated by a circular marker for each boxplot. For each region, the corresponding value of Cohen’s d is shown above the boxplot, to indicate effect size magnitude.

Overall, the two parameterizations produce qualitatively similar distribution spreads for most regions. However, the region-wise parameterization consistently predicts larger displacements, reflected by higher medians, means, and upper quartiles. Across both parameterizations, the largest displacements occur in the cerebellum, corona radiata, and cortex, with average values around 4 mm. In contrast, the corpus callosum, hippocampus, midbrain, and amygdala exhibit the smallest displacements, averaging around 2 mm.

These trends are not governed solely by regional material stiffness. While softer tissue generally exhibits larger deformation and stiffer tissue smaller deformation, the observed displacement field also contains contributions from rigid-body motion induced region interactions. As a result, local stiffness alone cannot fully explain the displacement magnitudes. Each region’s motion is influenced by the mechanical coupling with neighboring tissue properties, geometric location, and boundary constraints.

For example, the cerebellum, despite having an intermediate shear modulus, undergoes substantial displacement due to its contact with more compliant surrounding tissues and cerebrospinal fluid, allowing both deformation and bulk motion. Similarly, the corona radiata and cortex are located closer to the brain surface and interface with softer or fluid-filled regions, which allows larger overall motion. In contrast, deeply embedded regions such as the corpus callosum and hippocampus are surrounded by relatively stiffer tissues, including the cortex and basal ganglia, which restrict both deformation and rigid-body displacement. The midbrain and amygdala are further constrained by surrounding tissues and by the skull base through the homogeneous Dirichlet boundary condition.

Although most regions display generally similar displacement patterns between the two parameterization models, differences between them remain statistically significant, as supported by Cohen’s *d* values. The brain stem is the only region showing agreement between both parameterizations. They both predict wide displacement ranges, consistent with the broad spread of shear modulus values in this region (0–300 Pa voxel-wise and 100–450 Pa region-wise). Despite these differences in stiffness ranges, the resulting displacement fields are comparable, suggesting that regional heterogeneity averages out within the brain stem.

By contrast, the basal ganglia exhibit the largest discrepancy between the two parameterizations, with notable differences in the predicted displacements, their mean, and median. Interestingly, this occurs despite good agreement in the shear modulus distributions of the two parameterizations for the basal ganglia. The mismatch can be attributed to the mechanical environment: the basal ganglia are surrounded by regions with stiffnesses that differ between the parameterizations, including the hippocampus, for which the FA-based parameterization assigns higher stiffness.

#### . Principal stretches

Figure 10 shows the distributions of the first principal stretch across all regions for both parameterizations. Overall, the first principal stretch ranges between approximately 0.9 and 1.25. The corpus callosum experiences the highest tension, with an average tensile strain of about 5%, followed by the hippocampus, where tensile strains range between 1% and 10%. Most other regions undergo only mild tensile or compressive deformation, typically below 1%. This behavior is expected due to the intrinsic tension–compression asymmetry of brain tissue, which is captured by the one-term Ogden model. Across regions, the boxplots for the two material parameterizations align closely, indicating that the predicted deformations are similar with only small differences between the material parameterizations.

**Figure 10:**
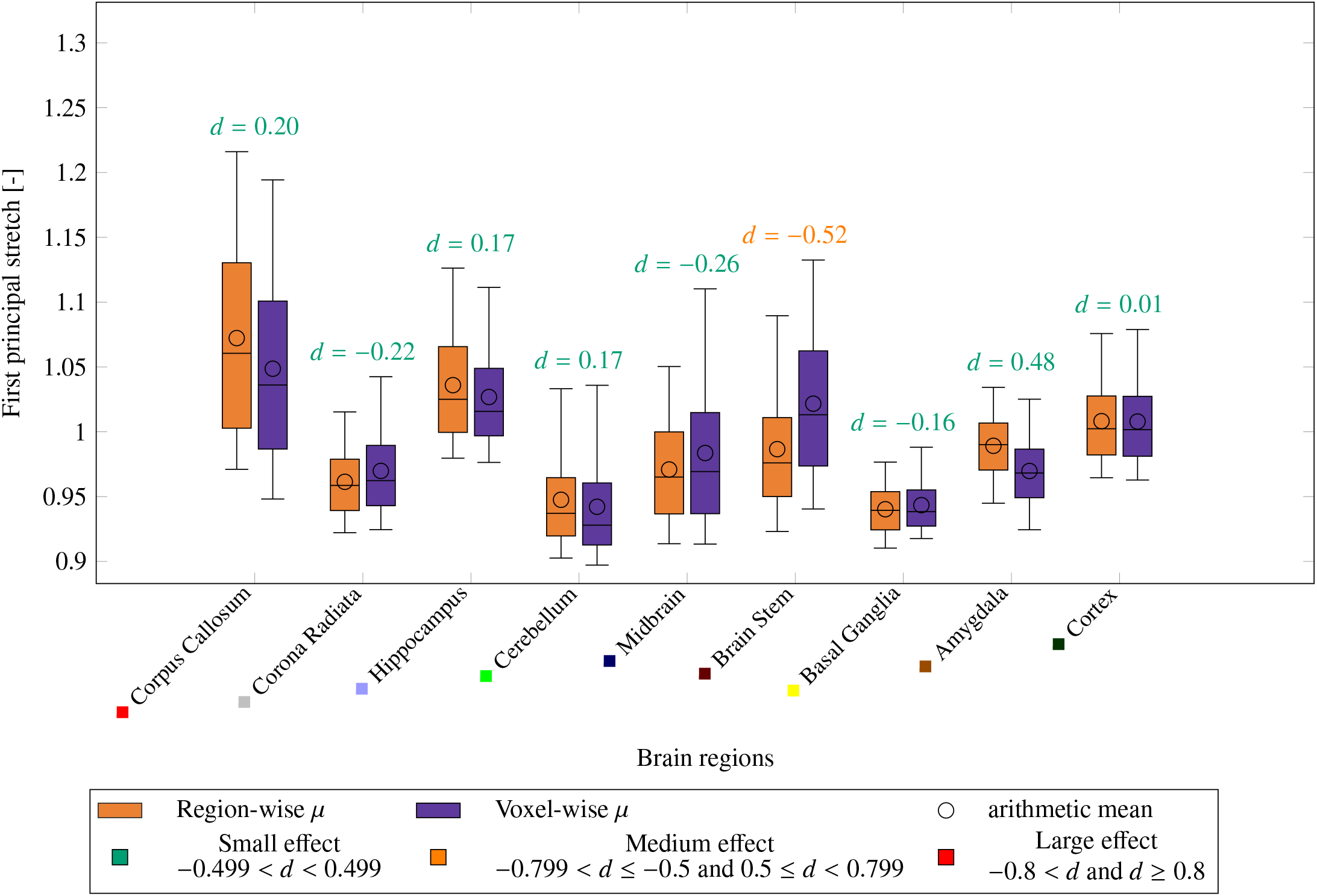
Boxplots illustrating the distributions of the simulated first principal stretch [-] across the nine anatomical brain regions for both the region-wise and voxel-wise parameterizations. Whiskers extend to the 5^th^ and 95^th^ percentiles, while the boxes and the central line indicate the 25^th^, 50^th^ (median), and 75^th^ percentiles; the arithmetic mean is indicated by a circular marker for each boxplot. For each region, the corresponding value of Cohen’s d is shown above the boxplot, to indicate effect size magnitude.

Regions that have on average a lower shear modulus *µ* in the region-wise parameterization, such as the corpus callosum, hippocampus, and cerebellum, show larger stretches in the region-wise simulations, consistent with lower stiffness that result in larger deformations. Conversely, regions that have on average a higher *µ* in the region-wise parameterization (midbrain, brainstem, basal ganglia) experience lower first principal stretches relative to the voxel-wise parameterization. Overall, all regions show good agreement, with Cohen’s d values between –0.26 and 0.48, indicating non-significant differences between distributions. The brainstem is the only region displaying a medium effect size. This deviation is explained by the higher stiffness assigned in the region-wise parameterization (≈ 200 Pa versus an average of ≈ 100 Pa in the voxel-wise FA-based parameterization).

The third principal stretch (Figure 11), which reflects compressive deformation, ranges from approximately 0.75 to 0.95. The corpus callosum undergoes the strongest compression (around 20%), whereas the cortex shows the least compressive behavior, with values approaching 10%. Most other regions fall within 10–20% compression. The voxel-wise parameterization frequently predicts less compression in regions where a higher shear modulus is assigned, consistent with a stiffer mechanical response. Although the two parameterizations exhibit broadly similar distributions, medium effect sizes appear in the corpus callosum and amygdala. In the corpus callosum, the relatively high *µ* in the voxel-wise parameterization leads to reduced compression, whereas in the basal ganglia, lower *µ* in the voxel-wise parameterization increases compression.

**Figure 11:**
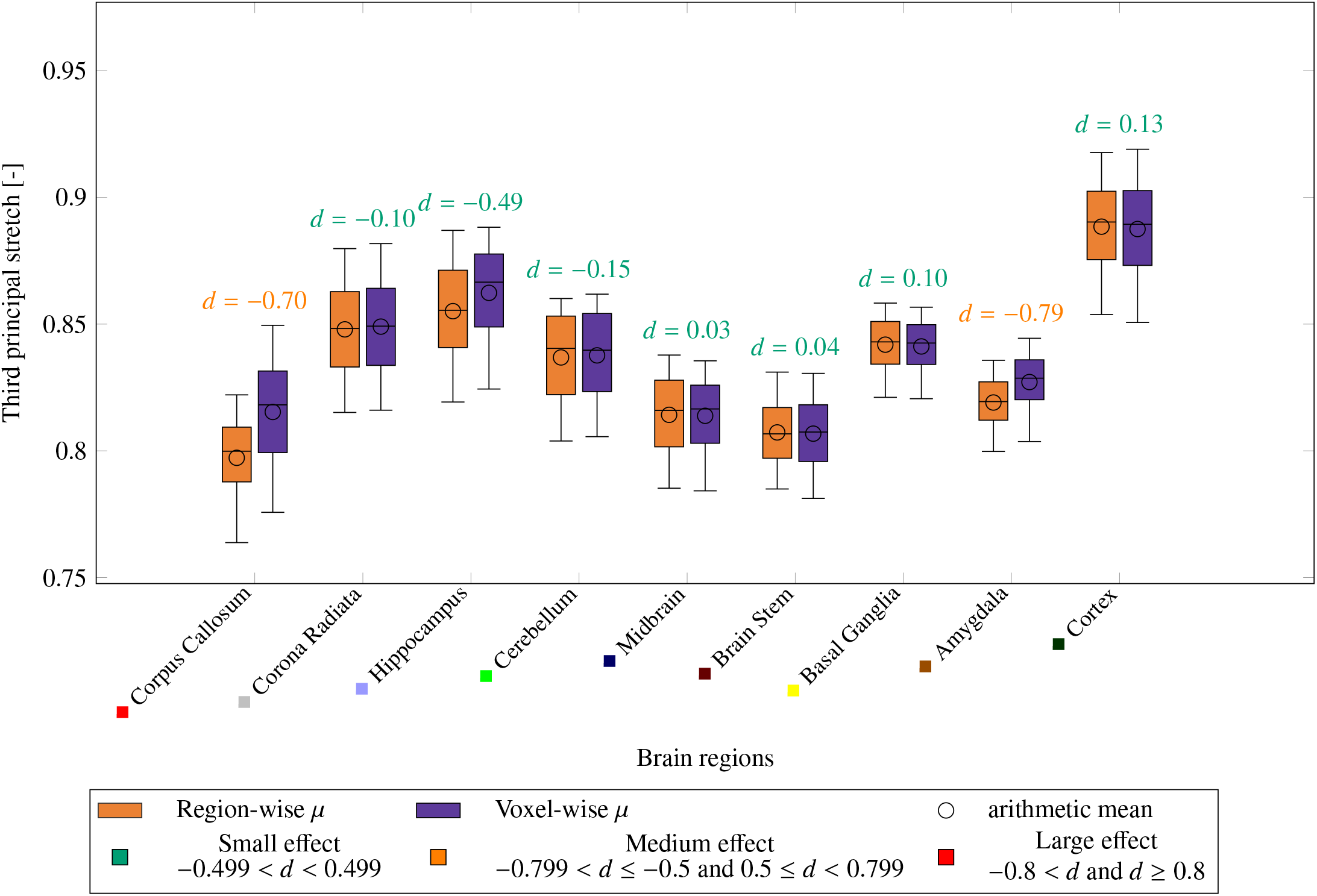
Boxplots illustrating the distributions of the simulated third principal stretch [-] across the nine anatomical brain regions for both the region-wise and voxel-wise parameterizations. Whiskers extend to the 5^th^ and 95^th^ percentiles, while the boxes and the central line indicate the 25^th^, 50^th^ (median), and 75^th^ percentiles; the arithmetic mean is indicated by a circular marker for each boxplot. For each region, the corresponding value of Cohen’s d is shown above the boxplot, to indicate effect size magnitude.

Finally, the maximum shear strain (Figure 12) ranges between 0 and 0.5 and reflects the difference between the first and third principal stretches. High values correspond to large deviations between tensile and compressive deformations, indicating greater distortion. The corpus callosum displays the highest shear in both parameterizations, consistent with its highly organized microstructure. This region contains densely packed, strongly aligned, predominantly left–right oriented fibers [41, 42, 43]. Such pronounced anisotropy enhances its susceptibility to both tensile and compressive deformation, increasing the difference between the first and third principal stretches and increasing shear. Moreover, diffusion MRI studies confirm the fiber orientation and dispersion across the corpus callosum [44, 45], which may further contribute to the broader shear distributions observed in both parameterizations. Regions with high spatial variability in the voxel-wise shear modulus, such as the midbrain and the brainstem, present more compact shear distributions in the voxel-wise simulations. Incorporating local stiffness heterogeneity disperses strain gradients and can reduce peak shear. In contrast, the region-wise parameterization assigns uniform stiffness within each anatomical region, which can allow higher localized shear concentrations.

**Figure 12:**
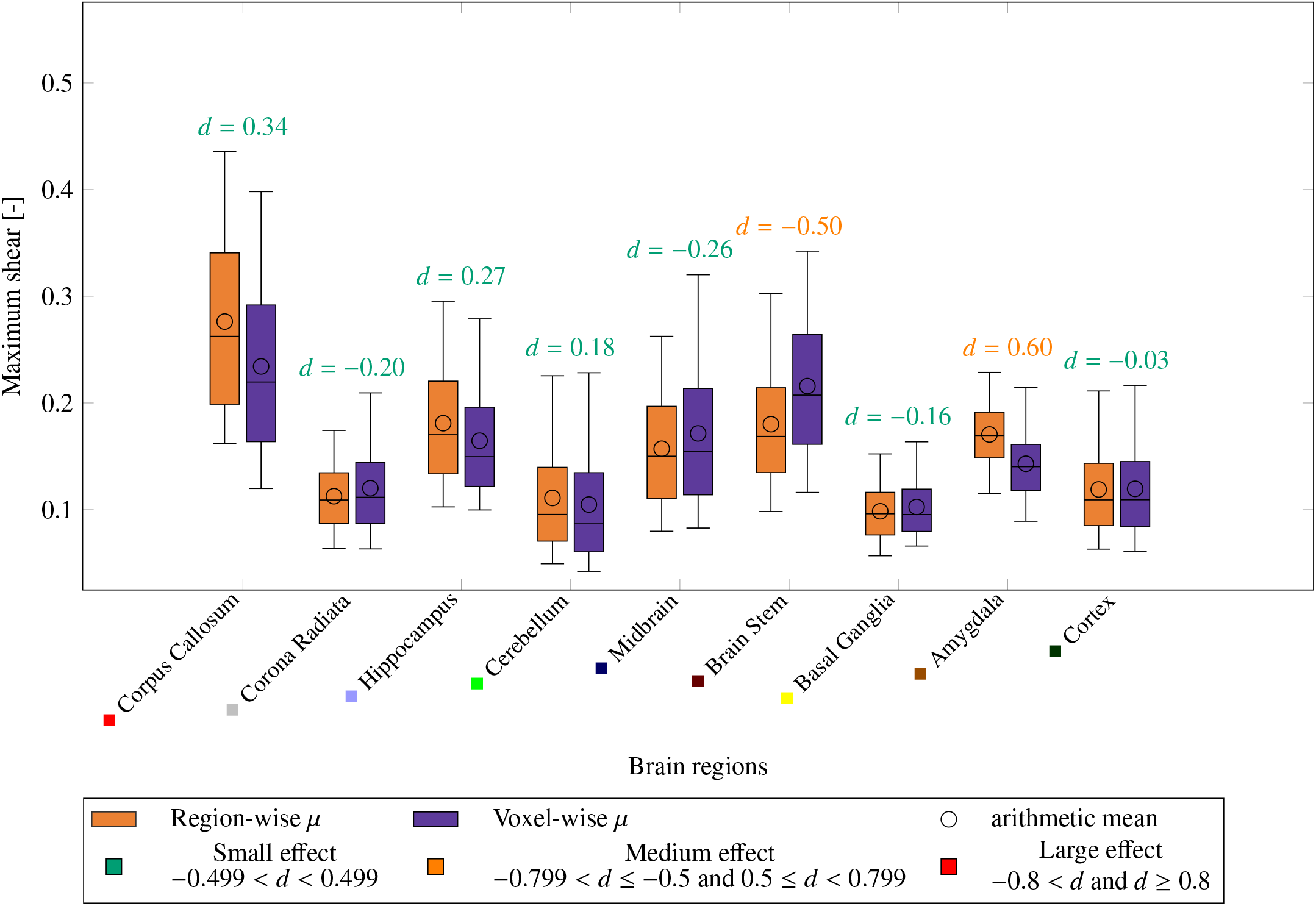
Boxplots illustrating the distribution of the maximum shear [-] across the nine anatomical brain regions for both the region-wise and voxel-wise parameterizations. Whiskers extend to the 5^th^ and 95^th^ percentiles, while the boxes and the central line indicate the 25^th^, 50^th^ (median), and 75^th^ percentiles; the arithmetic mean is indicated by a circular marker for each boxplot. For each region, the corresponding value of Cohen’s d is shown above the boxplot, to indicate effect size magnitude.

## Conclusion

In this work, we developed and compared two biomechanical parameterizations of the same human brain: (i) a region-wise parameterization with piecewise-constant material parameters, and (ii) a voxel-wise FA-based parameterization in which the shear modulus varies spatially according to local diffusion-derived microstructure. Using experimental 9-region mechanical data, we established a linear relationship between fractional anisotropy (FA) and shear modulus *µ*, which allowed to define a spatially heterogeneous stiffness field by applying this mapping voxel-wise. Both parameterizations were subjected to identical cerebral atrophy simulations, and the resulting deformation, stretches, and volumetric changes were compared.

A strong, statistically significant negative correlation was found between regional FA and shear modulus, supporting the use of FA as an indicator of local stiffness. Within the corpus callosum, high FA values produced negative predicted stiffness for some voxels, necessitating a cutoff. Interestingly, despite the applied FA threshold, the FA-derived *µ* values still aligned with the region-wise parameterization values for the same region.

The voxel-wise parameterization produced a continuous, microstructurally informed stiffness distribution with large variability within each region, unlike the piecewise constant stiffness per region observed in the 9R parameterization. Despite these differences, both parameterizations predicted similar global atrophy. However, the voxel-wise parameterization predicted greater ventricular enlargement, due to softer periventricular tissue revealed by the FA-derived shear modulus. Locally, the mechanical responses diverged. Most regions displayed revealed differences in displacement. These discrepancies stem from the broad *µ* distributions within and surrounding these regions, that the region-wise parameterization cannot capture. The corpus callosum also exhibited a wide distribution of shear strain, likely reflecting its highly ordered left-right fiber architecture.

This study introduces several methodological advances. First, we demonstrate a direct voxel-wise stiffness derived from FA using experimentally identified region-level mechanics. Second, the approach provides a fully non-invasive means to obtain patient-specific, spatially varying material properties for brain simulations, based on existing experimental data. Finally, our findings illustrate that microstructural heterogeneity can meaningfully influence local mechanical responses.

The study has several limitations. The FA-*µ* mapping relies on a simple linear regression, which may not capture potential nonlinear or region-specific microstructure-mechanics relationships. The nonlinearity parameter *α* was assumed constant within each region due to the absence of FA correlations, which may underestimate differences in nonlinear behavior. The constitutive model remains isotropic, despite well-known anisotropic features of white matter. Atrophy was modeled as isotropic shrinkage-appropriate for comparing the two material parameterizations, though not for a comprehensive representation of disease processes. Finally, and most importantly, determining which parameterization better reflects in vivo brain biomechanics will require dedicated validation in future studies.

## Data Availability

The datasets generated in this study, including the simulation outputs for both region-wise and FA-based parameterizations, are publicly available on Zenodo (https://doi.org/10.5281/zenodo.19571279).

## Acknowledgements

We further acknowledge the financial support by the Deutsche Forschungsgemeinschaft (DFG, German Research Foundation) through the grant BU 3728/1-1 and project 460333672 CRC1540 EBM (project A02).

## Appendix A. 3D Model

## Appendix B. Histograms

## Appendix C. Linear fits

## Appendix D. Mechanical outputs

**Figure A.13:**
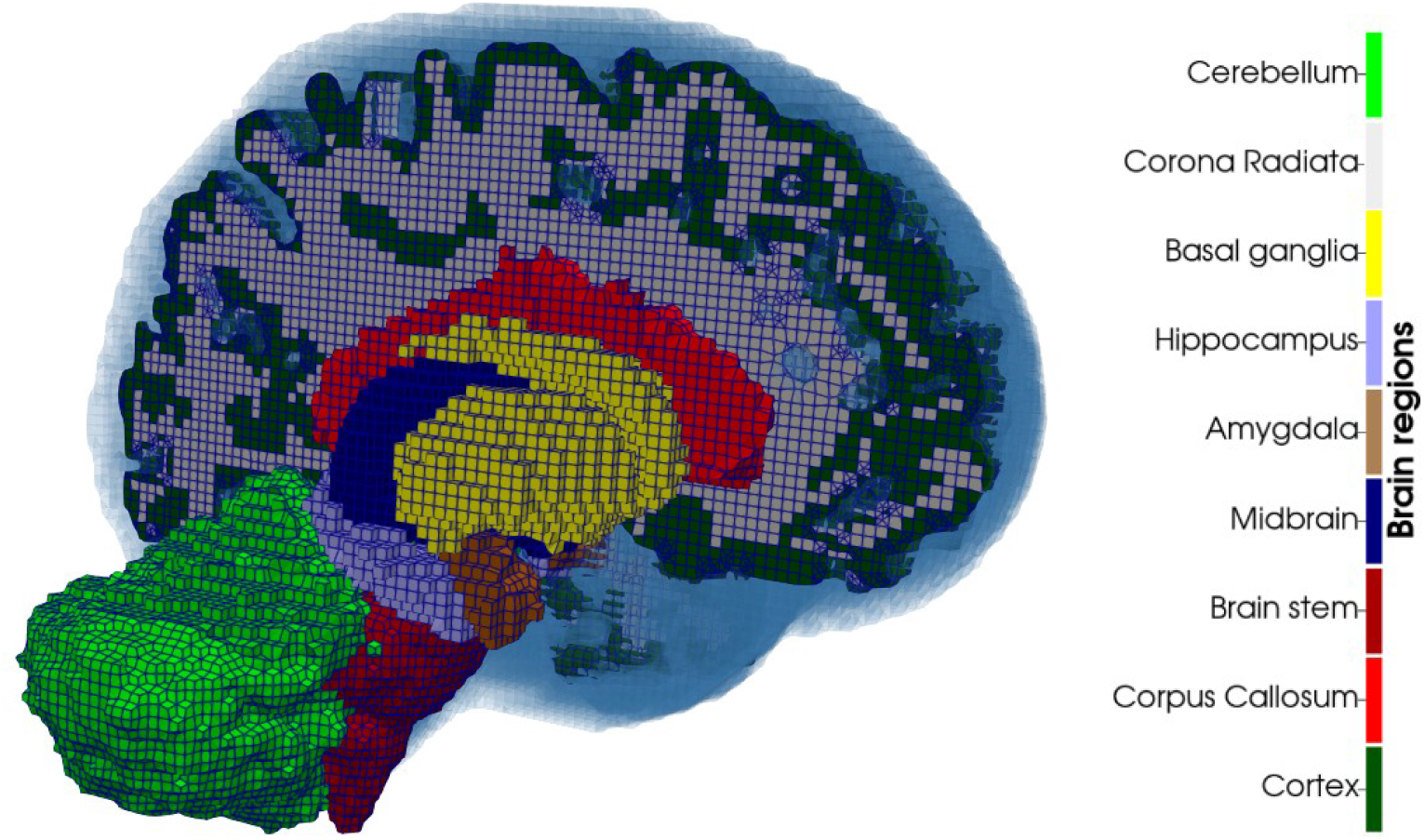
Regional segmentation for the 9-Region parameterization. The surrounding cerebrospinal fluid (CSF) layer is shown in blue.

**Table C.4:** Overview of the different linear fit functions for the four *α* value datasets and the corresponding statistics of the Pearson test.

| Dataset | Linear mapping | $R^2$ | $p_r$ |
| --- | --- | --- | --- |
| Preconditioned, $\nu = 0.45$ | $v_{FA} = -0.0056 \alpha + 0.2001$ | 0.0056 | 0.8484 |
| Preconditioned, $\nu = 0.49$ | $v_{FA} = -0.0105 \alpha + 0.1334$ | 0.0898 | 0.4333 |
| Unconditioned, $\nu = 0.45$ | $v_{FA} = -0.0102 \alpha + 0.1338$ | 0.0362 | 0.6239 |
| Unconditioned, $\nu = 0.49$ | $v_{FA} = -0.0183 \alpha + 0.0902$ | 0.0637 | 0.5124 |

**Figure B.14:**
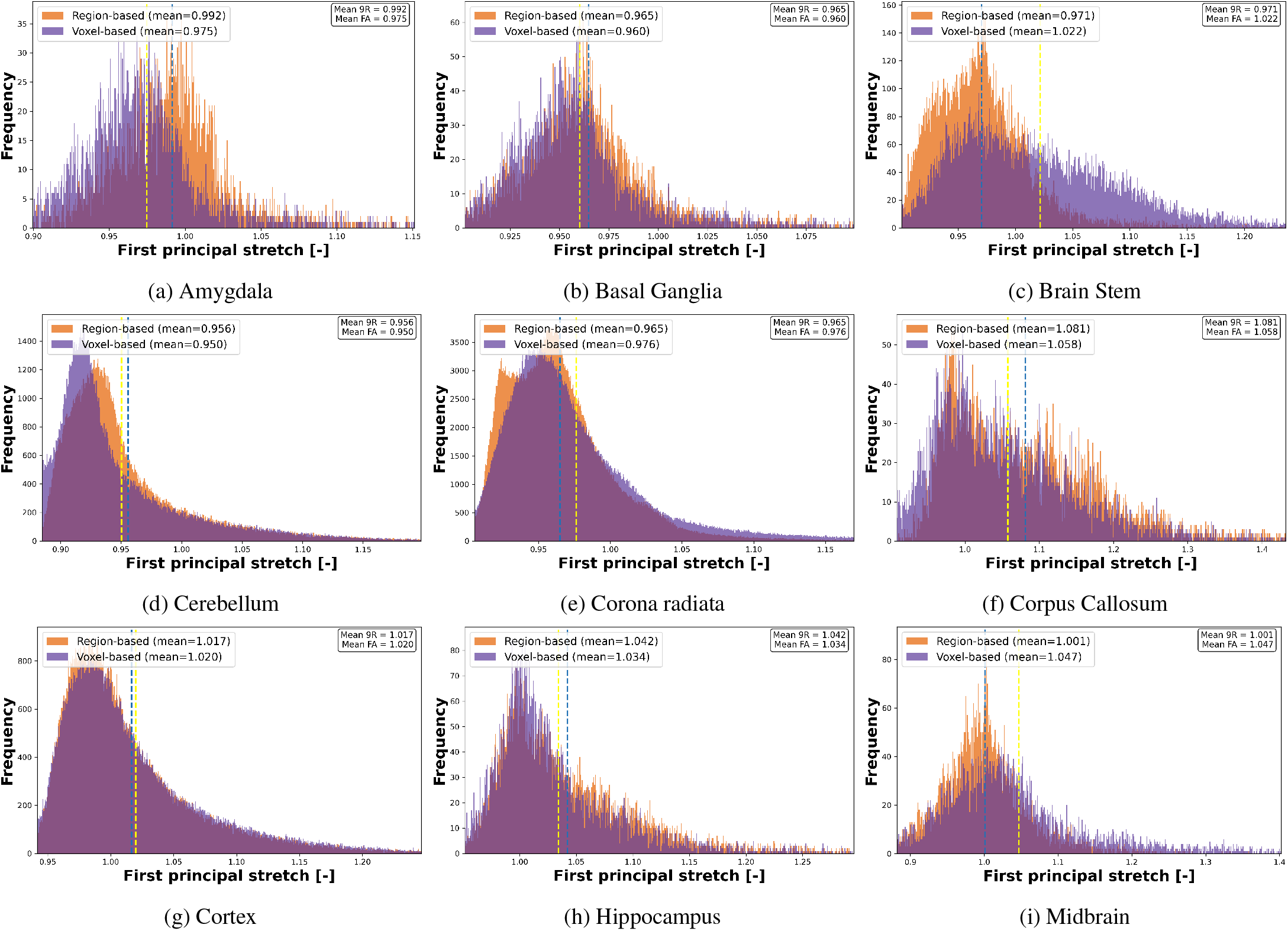
Histograms of the first principal stretch distribution across all brain regions for the region-wise (blue) and voxel-wise FA-based (yellow) parameterizations. The mean values for each parameterization are indicated by dashed lines in their respective colors. Data were cropped to the 1st–99th percentile range prior to plotting to reduce the impact of extreme outliers.

**Figure B.15:**
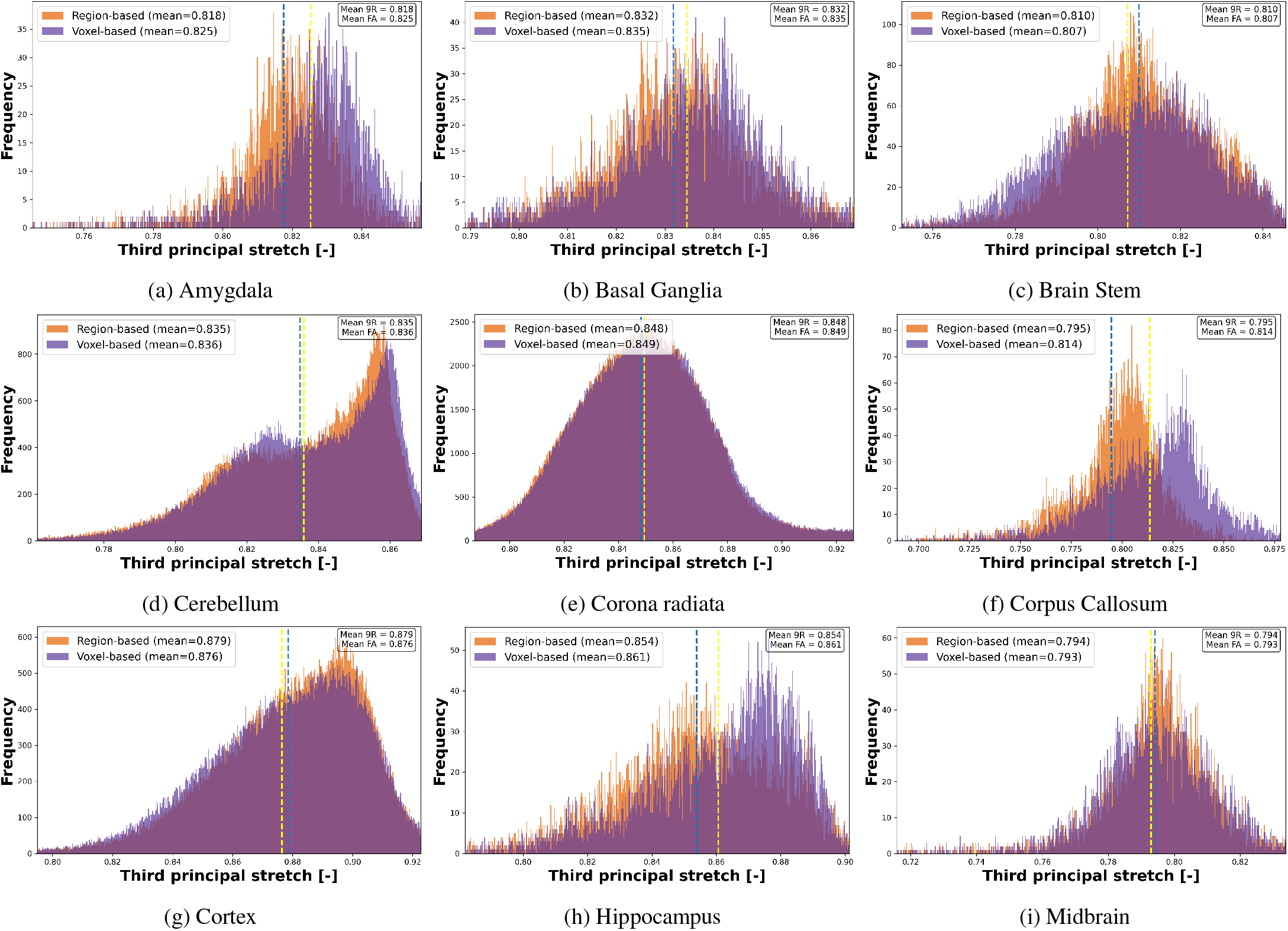
Histograms of the third principal stretch distribution across all brain regions for the region-wise (blue) and voxel-wise FA-based (yellow) parameterizations. The mean values for each parameterization are indicated by dashed lines in their respective colors. Data were cropped to the 1st–99th percentile range prior to plotting to reduce the impact of extreme outliers.

**Figure B.16:**
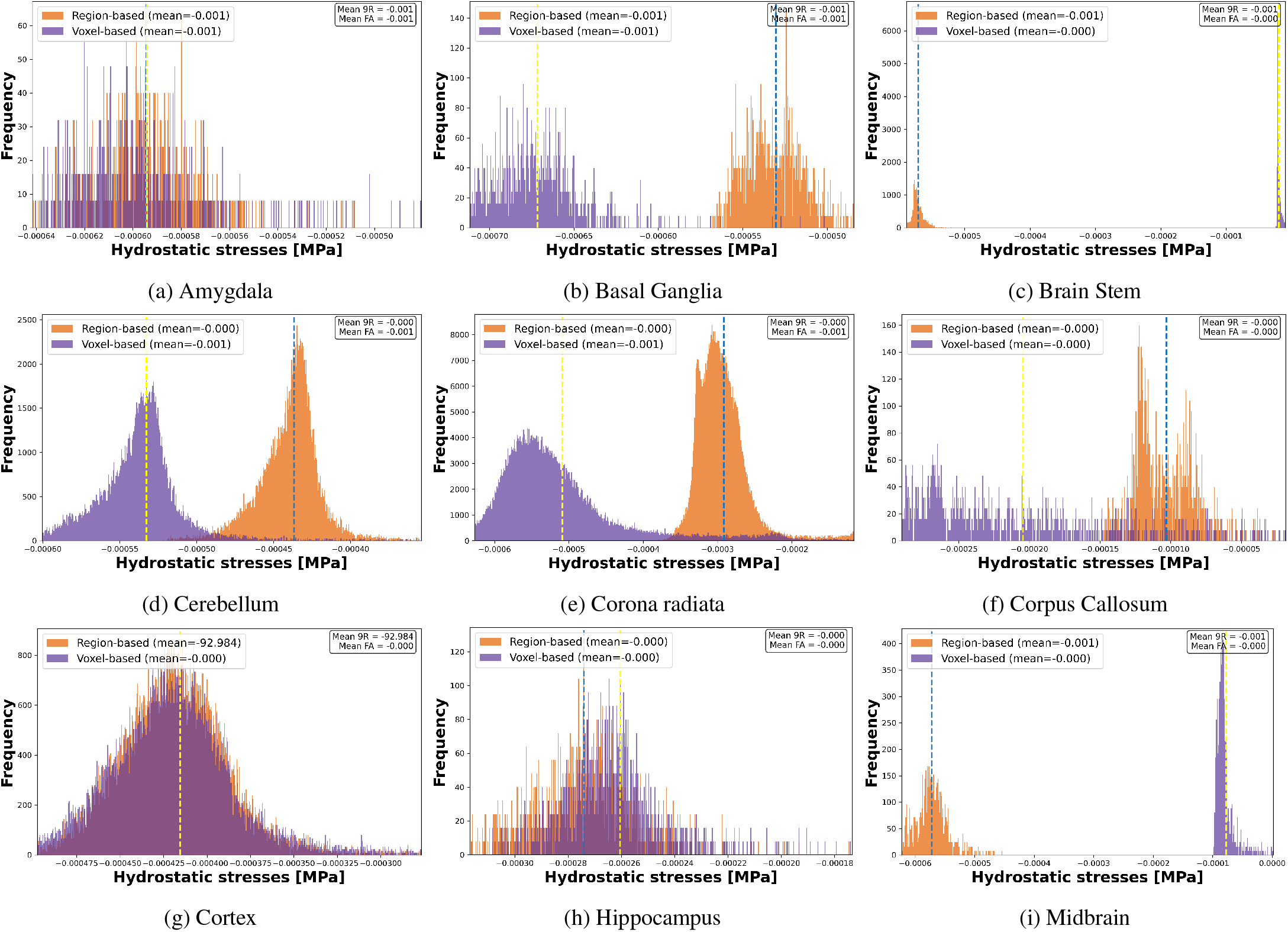
Histograms of the hydrostatic stress distribution across all brain regions for the region-wise (blue) and voxel-wise FA-based (yellow) parameterizations. The mean values for each parameterization are indicated by dashed lines in their respective colors. Data were cropped to the 1st–99th percentile range prior to plotting to reduce the impact of extreme outliers.

**Figure B.17:**
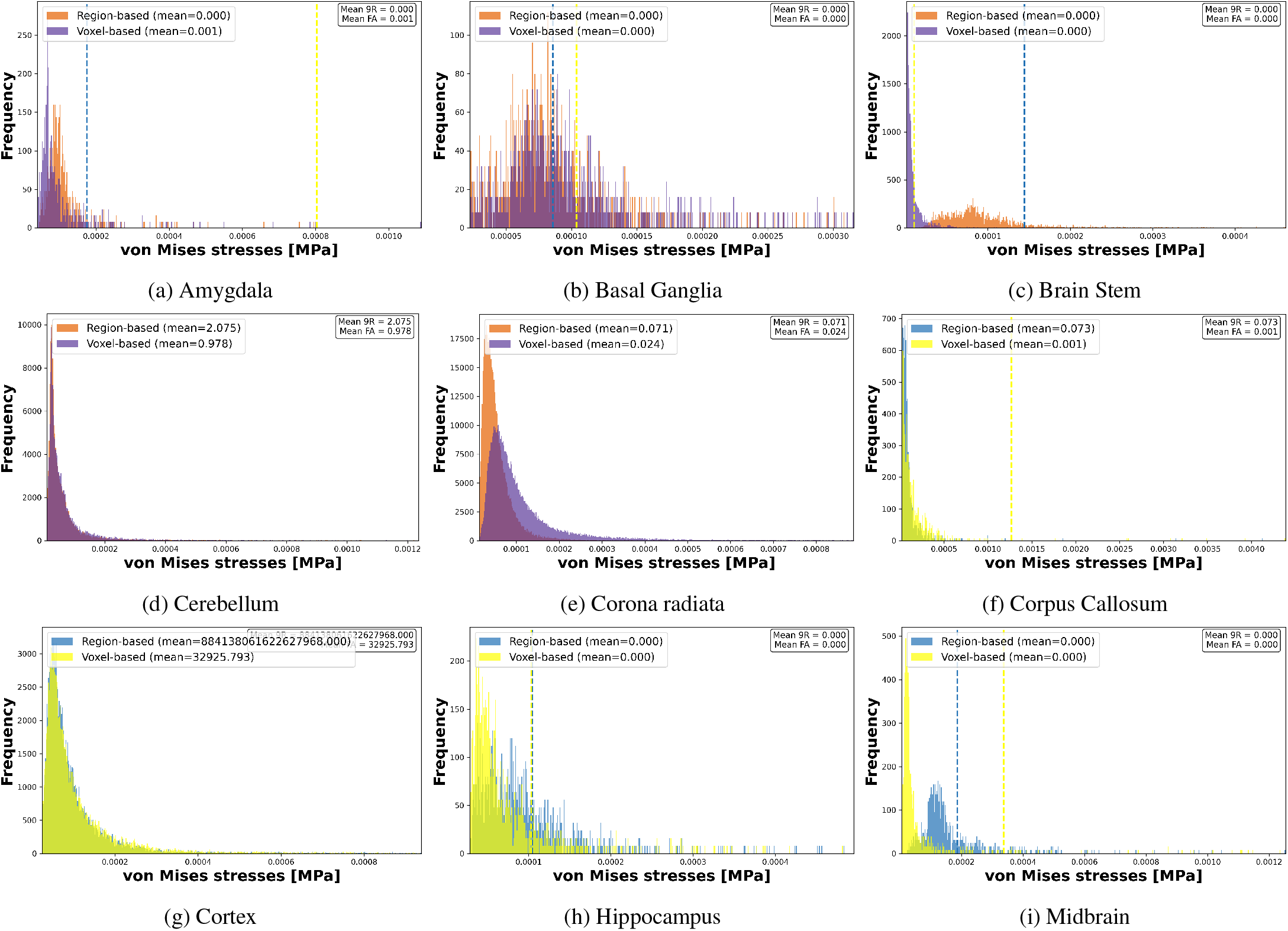
Histograms of the von-Mises stress distribution across all brain regions for the region-wise (blue) and voxel-wise FA-based (yellow) parameterizations. The mean values for each parameterization are indicated by dashed lines in their respective colors. Data were cropped to the 1st–99th percentile range prior to plotting to reduce the impact of extreme outliers.

**Figure C.18:**
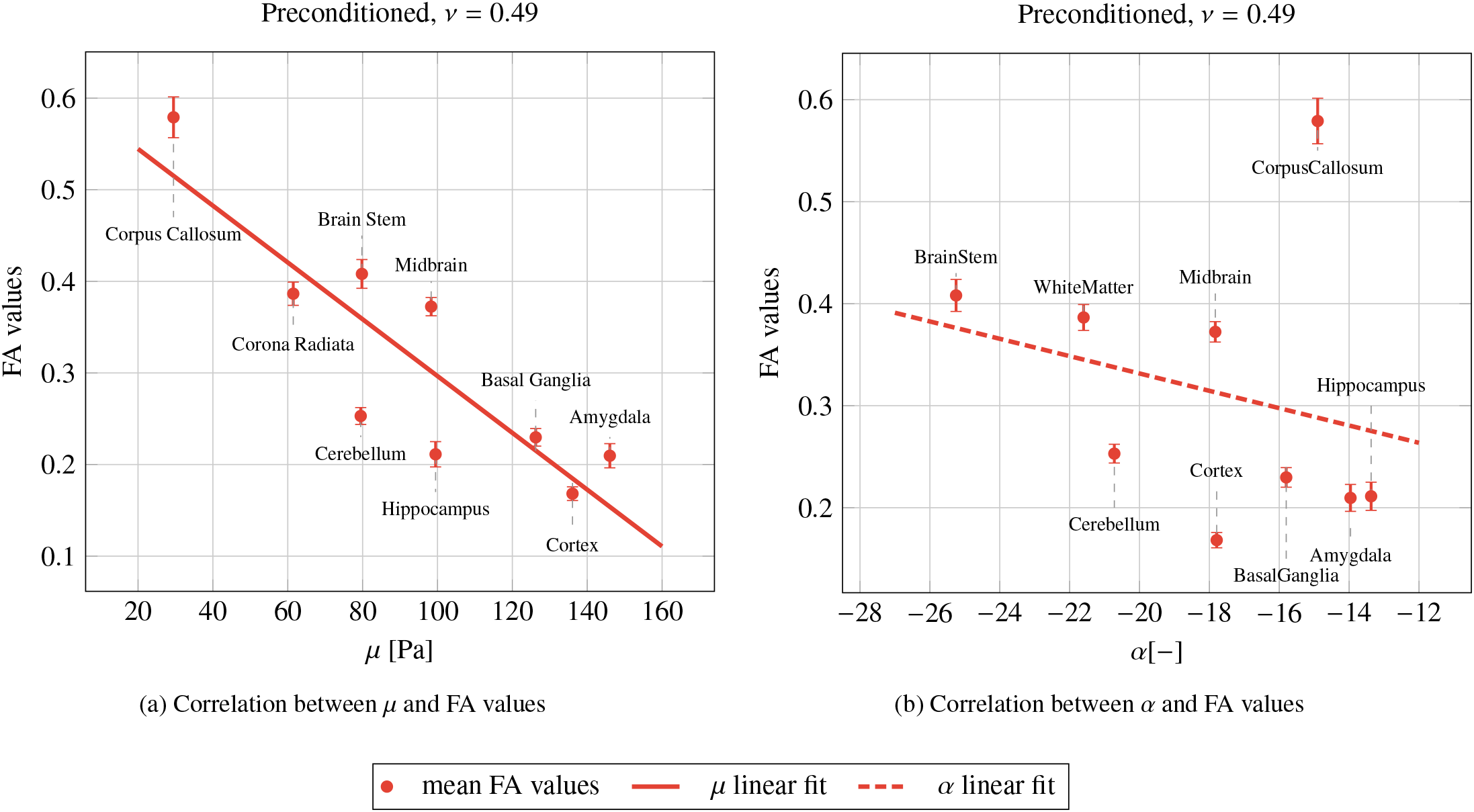
Comparison between *µ* (left) and *α* (right) correlations for the preconditioned, *ν* = 0.49 dataset.

**Figure C.19:**
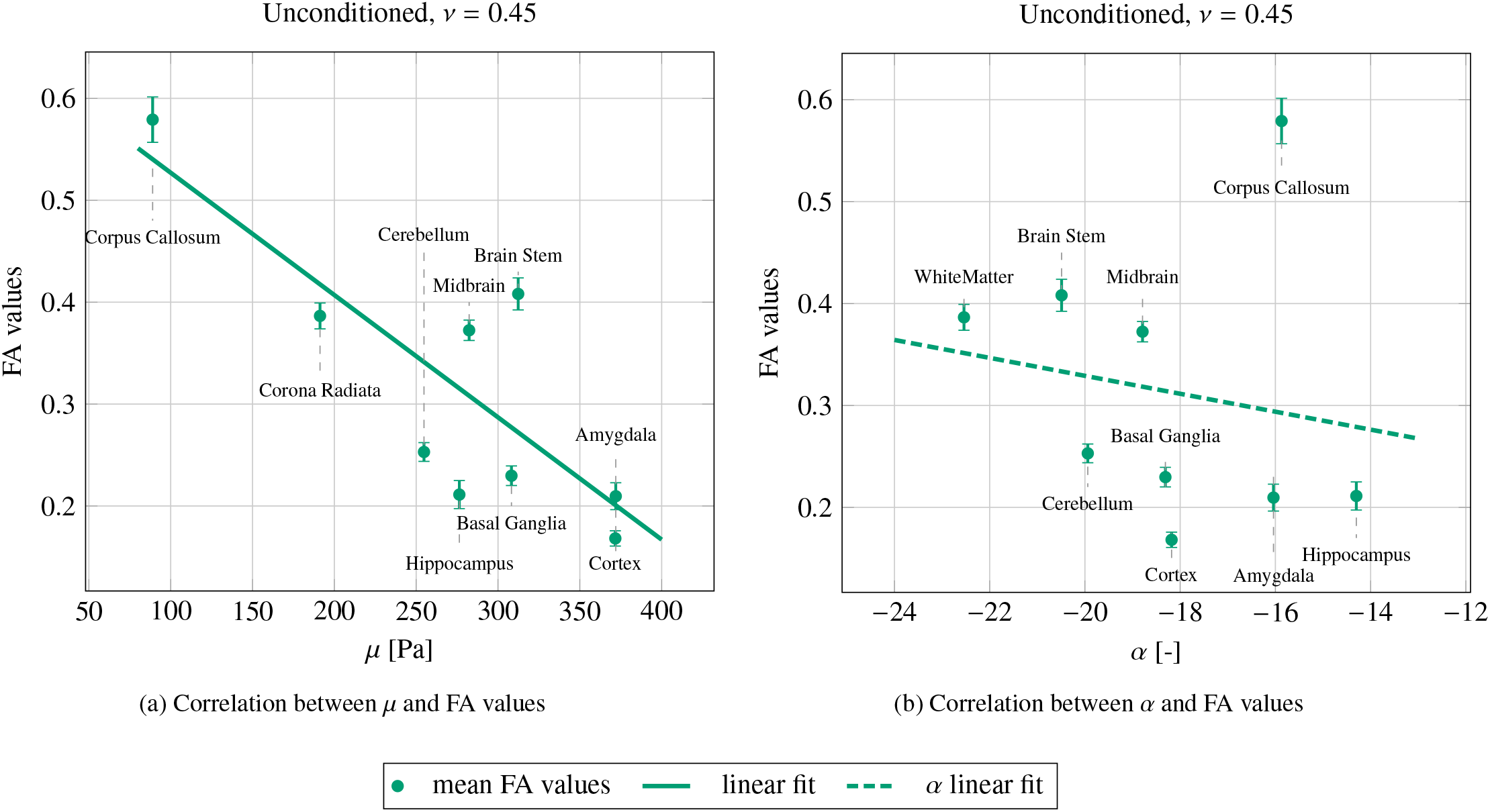
Comparison between *µ* (left) and *α* (right) correlations for the unconditioned, *ν* = 0.45 dataset.

**Figure C.20:**
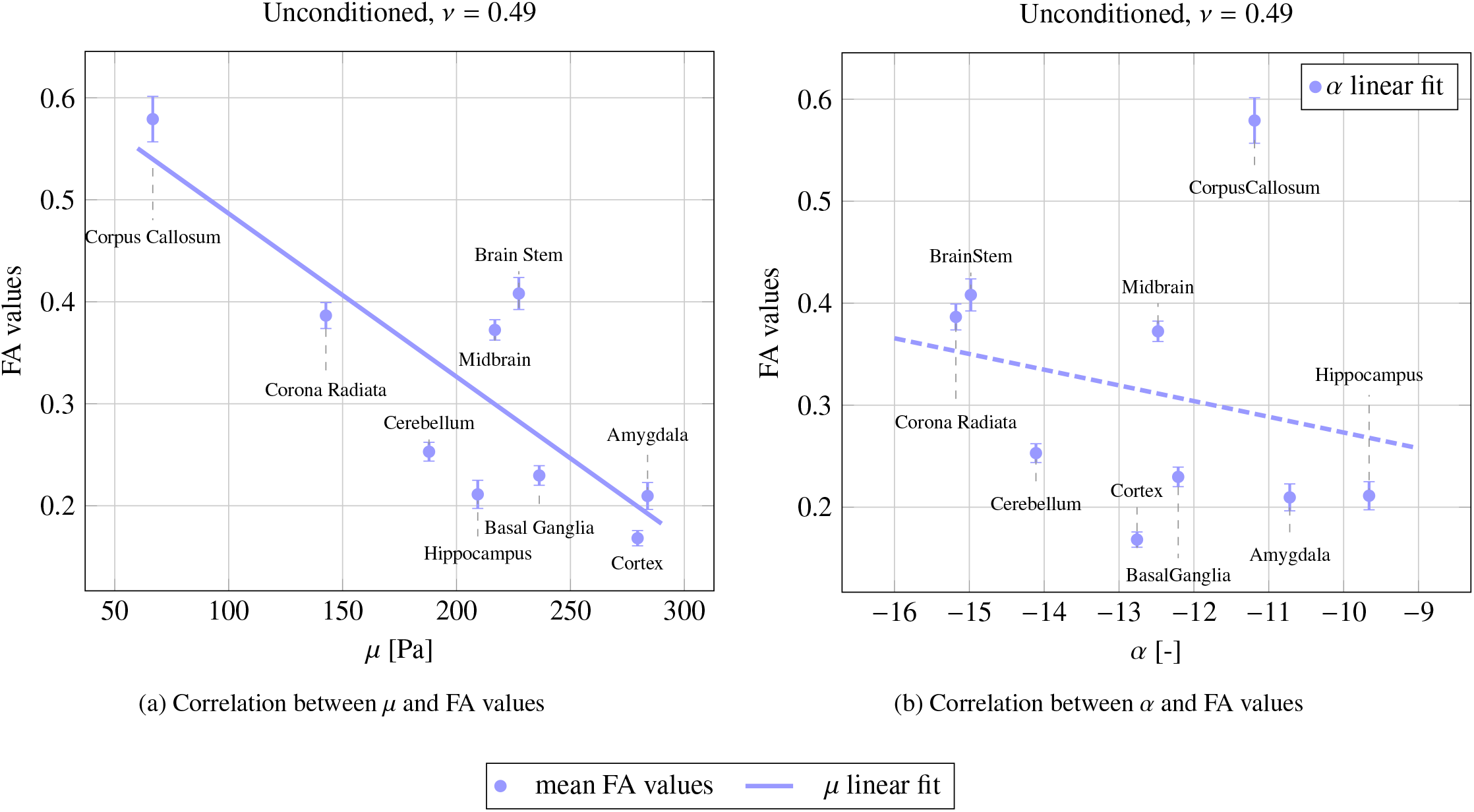
Comparison between *µ* (left) and *α* (right) correlations for the unconditioned, *ν* = 0.49 dataset.

**Figure D.21:**
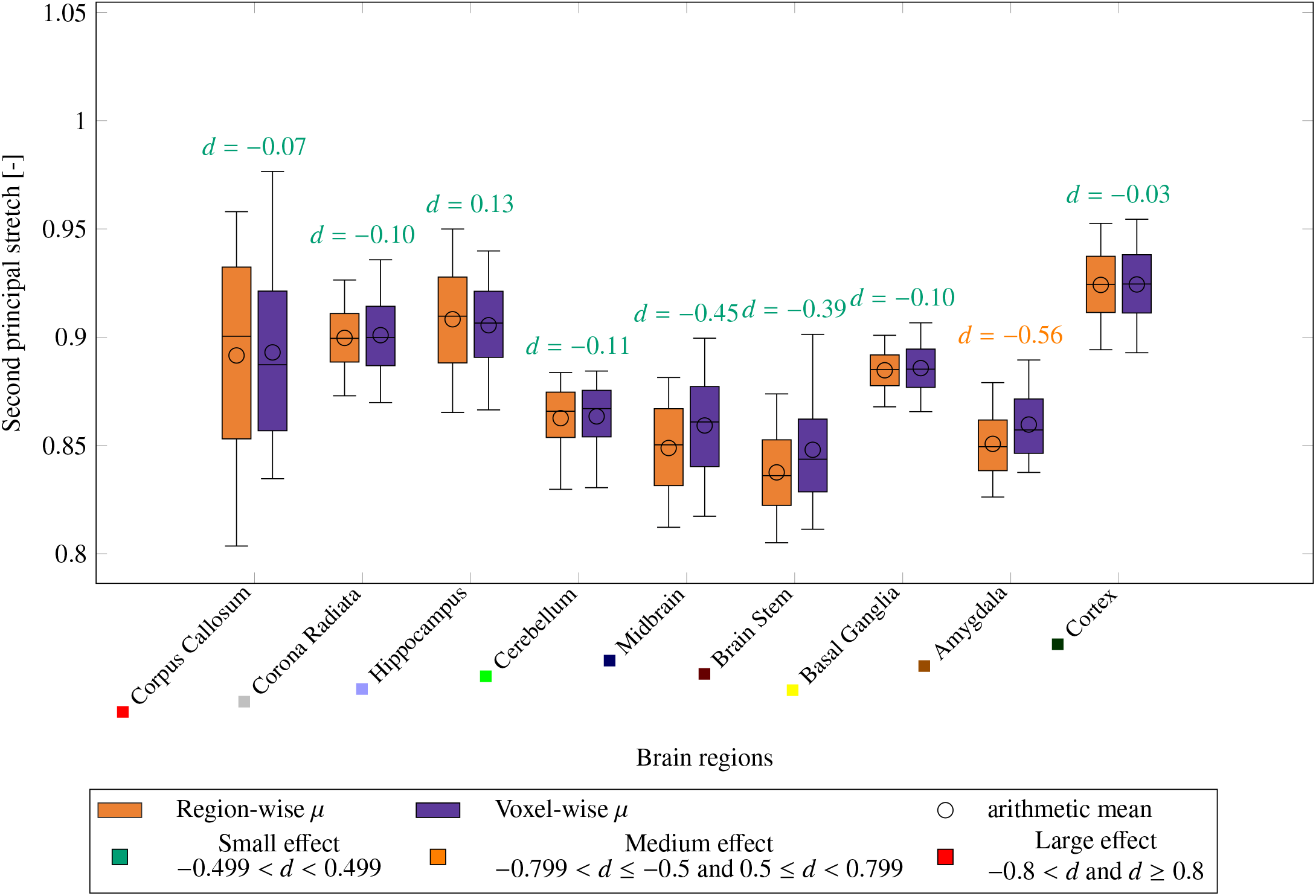
Boxplots illustrating the distributions of the simulated second principal stretch [-] across the nine anatomical brain regions for both the region-wise and voxel-wise parameterizations. Whiskers extend to the 5^th^ and 95^th^ percentiles, while the boxes and the central line indicate the 25^th^, 50^th^ (median), and 75^th^ percentiles; the arithmetic mean is indicated by a circular marker for each boxplot. For each region, the corresponding value of Cohen’s d is shown above the boxplot, to indicate effect size magnitude.

**Figure D.22:**
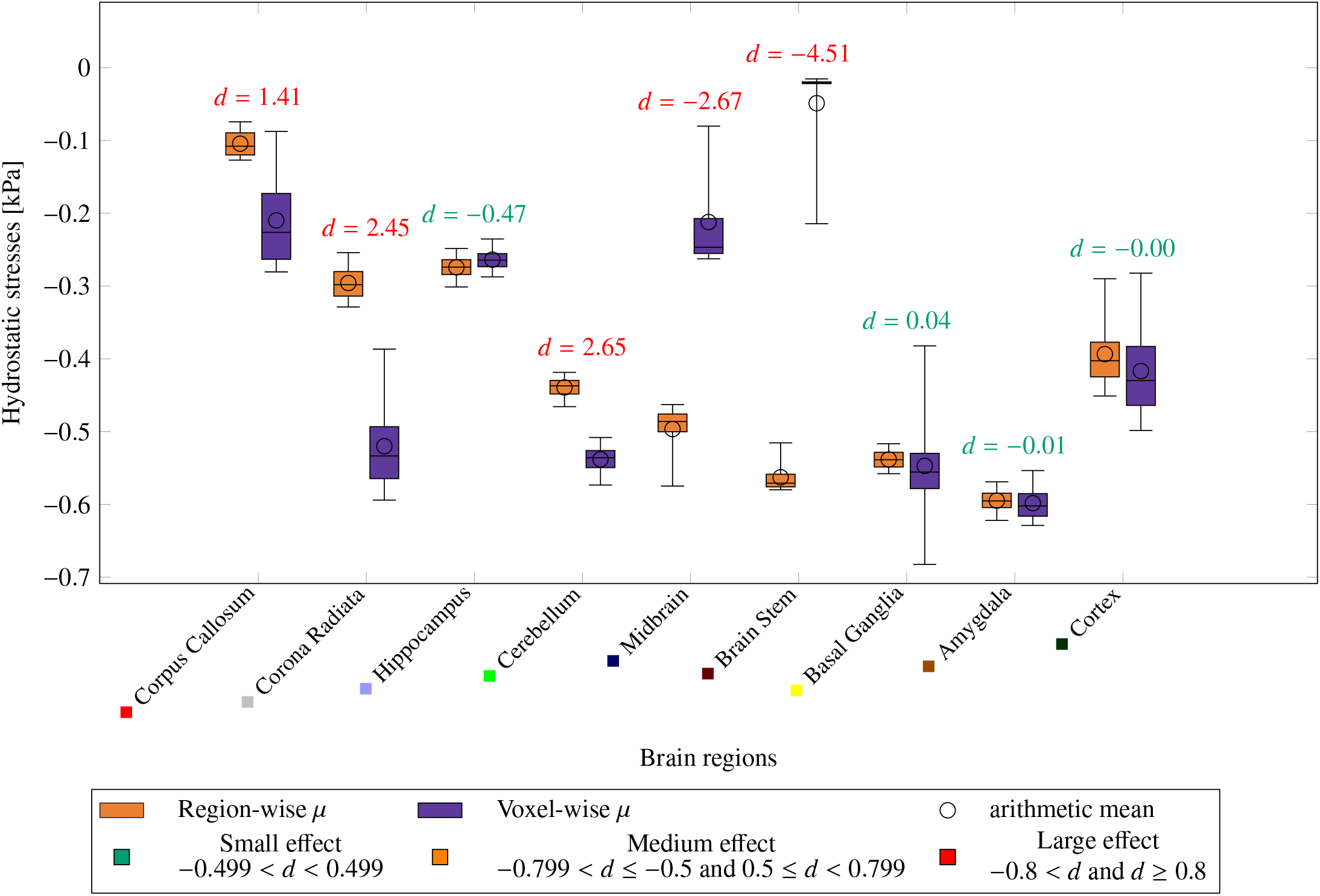
Boxplots illustrating the distributions of the hydrostatic stresses [kPa] across the nine anatomical brain regions for both the region-wise and voxel-wise parameterizations. Whiskers extend to the 5^th^ and 95^th^ percentiles, while the boxes and the central line indicate the 25^th^, 50^th^ (median), and 75^th^ percentiles; the arithmetic mean is indicated by a circular marker for each boxplot. For each region, the corresponding value of Cohen’s d is shown above the boxplot, to indicate effect size magnitude.

**Figure D.23:**
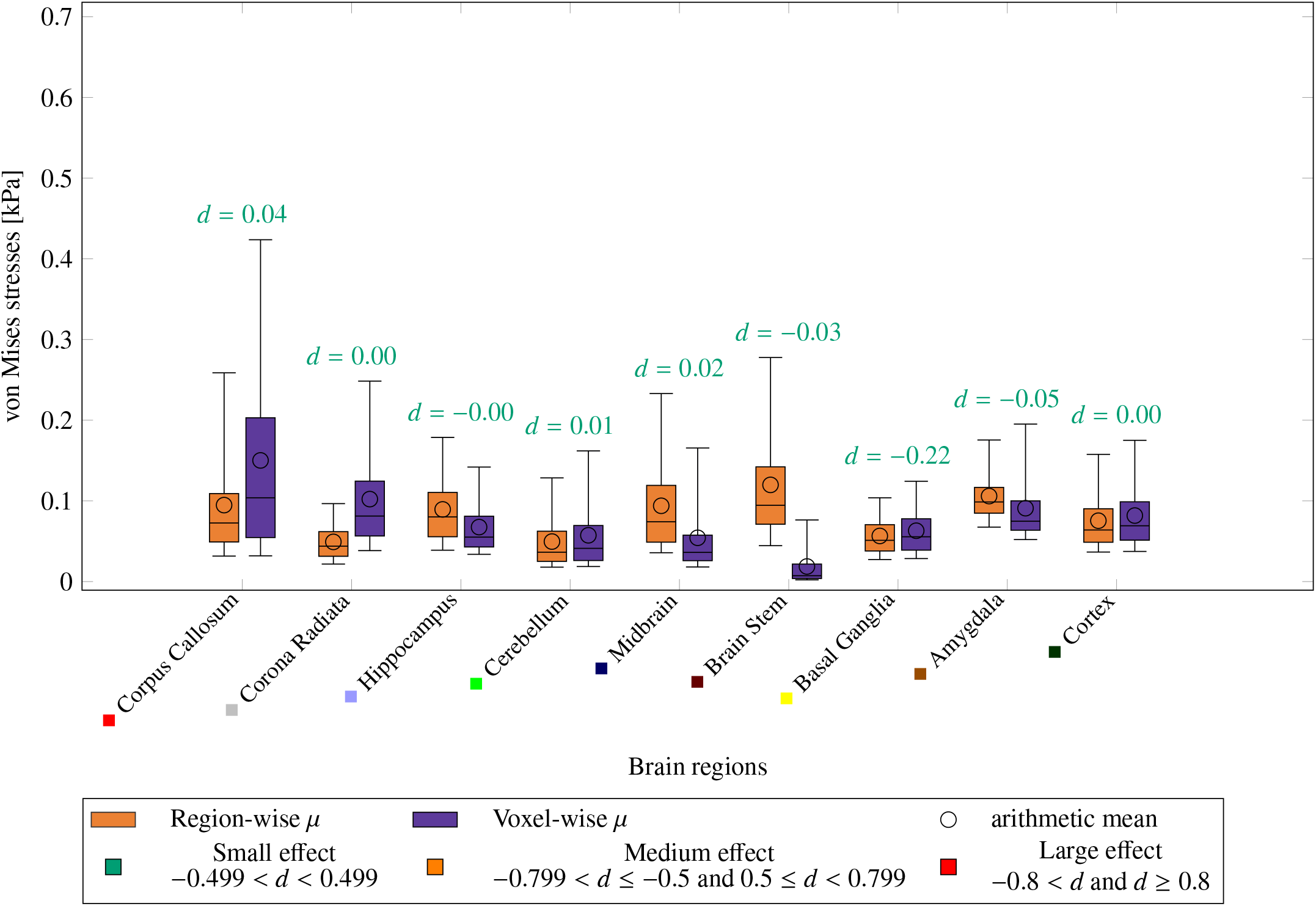
Boxplots illustrating the distributions of the simulated von Mises stresses [kPa] across the nine anatomical brain regions for both the region-wise and voxel-wise parameterizations. Whiskers extend to the 5^th^ and 95^th^ percentiles, while the boxes and the central line indicate the 25^th^, 50^th^ (median), and 75^th^ percentiles; the arithmetic mean is indicated by a circular marker for each boxplot. For each region, the corresponding value of Cohen’s d is shown above the boxplot, to indicate effect size magnitude.

## Footnotes

1 https://surfer.nmr.mgh.harvard.edu/

2 https://github.com/BRAINIACS-Group/Brain-Creation-Code/tree/FA_integration

3 https://github.com/BRAINIACS-Group/EFI_atrophy_and_tumor/tree/atrophy_FA_integration

